# Pulsatile lipid vesicles under osmotic stress

**DOI:** 10.1101/070342

**Authors:** Morgan Chabanon, James C.S. Ho, Bo Liedberg, Atul N. Parikh, Padmini Rangamani

**Affiliations:** Department of Mechanical and Aerospace Engineering, University of California San Diego, La Jolla, CA, USA; Center for Biomimetic Sensor Science, School of Materials Science and Engineering, Nanyang Technological University, Singapore; Departments of Applied Science, Biomedical Engineering, and Chemical Engineering and Materials Science, University of California Davis, Davis, CA, USA

## Abstract

The response of lipid bilayers to osmotic stress is an important part of cellular function. Previously, in (Oglecka et al., 2014), we reported that cell-sized giant unilamellar vesicles (GUVs) exposed to hypotonic media, respond to the osmotic assault by undergoing a cyclical sequence of swelling and bursting events, coupled to the membrane’s compositional degrees of freedom. Here, we seek to deepen our quantitative understanding of the essential pulsatile behavior of GUVs under hypotonic conditions, by advancing a comprehensive theoretical model for vesicle dynamics. The model quantitatively captures our experimentally measured swell-burst parameters for single-component GUVs, and reveals that thermal fluctuations enable rate dependent pore nucleation, driving the dynamics of the swell-burst cycles. We further identify new scaling relationships between the pulsatile dynamics and GUV properties. Our findings provide a fundamental framework that has the potential to guide future investigations on the non-equilibrium dynamics of vesicles under osmotic stress.

---

In their constant struggle with the environment, living cells of contemporary organisms employ a variety of highly sophisticated molecular mechanisms to deal with sudden changes in their surroundings. One often encountered environmental assault on cells is that of osmotic stress, where the amount of dissolved molecules in the extracellular environment drops suddenly (Christensen, 1987; Hoffmann et al., 2009). If unchecked, this perturbation will result in a rapid flow of water into the cell through osmosis, causing it to swell, rupture, and die. To avoid this catastrophic outcome, even bacteria have evolved complex molecular machineries, such as mechanosensitive channel proteins, which allow them to release excess water from their interior (Berrier et al., 1996; Blount et al., 1997; Levina et al., 1999; Wood, 1999). This then raises an intriguing question of how might primitive cells, or cell-like artificial constructs, that lack a sophisticated protein machinery, respond to such environmental insults to preserve their structural integrity.

Using rudimentary cell-sized giant unilamellar vesicles (GUVs) devoid of proteins and consisting of amphiphilic lipids and cholesterol as models for simple protocells, we showed previously in (Oglecka et al., 2014) that vesicular compartments respond to osmotic assault created by the exposure to hypotonic media, by undergoing a cyclical sequence of swelling and poration. In each cycle, osmotic influx of water through the semi-permeable boundary swells the vesicles and renders the bounding membrane tense, which in turn, opens a microscopic transient pore, releasing some of the internal solutes before resealing. This swell-burst process repeats multiple times producing a pulsating pattern in the size of the vesicle undergoing osmotic relaxation. From a dynamical point of view, this autonomous osmotic response results from an initial, far-from-equilibrium, thermodynamically unstable state generated by the sudden application of osmotic stress.

The subsequent evolution of the system, characterized by the swell-burst sequences described above, occurs in the presence of a global constraint, namely the constant membrane area, during a dissipation-dominated process (Peterlin and Arrigler, 2008; Ho et al., 2016). While we provided a qualitative interpretation of the vesicles pulsatile behavior (see (Oglecka et al., 2014) Fig. 7h,i), a quantitative description of the long time scale dynamics of swell-burst cycles is still needed to obtain insight into the underlying physics. Here, we build directly on the qualitative findings we previously reported to propose such a quantitative and mechanistic analysis of the dynamics of swell-burst cycles in lipid vesicles undergoing osmotic stress.

In analyzing the pulsatile dynamics of GUVs, a number of general questions naturally arise: (i) Is the observed condition for membrane poration deterministic or stochastic? (ii) Is poration controlled by a unique value of membrane tension (*i.e.* lytic tension) introduced by the area-volume changes, which occur during osmotic influx, or does it involve coupling the membrane response to thermal fluctuations? (iii) Does the critical lytic tension depend on the strain rate, and thus the strength of the osmotic gradient? Such questions arise, beyond the present context of vesicle osmoregulation, in other important scenarios where the coupling between the dissipation of osmotic energy and cellular compartmentalization have important biological ramifications (Rand, 2004; Diz-Muñoz et al., 2013; Stroka et al., 2014; Porta et al., 2015).

Motivated by these considerations, we carried out a combined theoretical-experimental study integrating membrane elasticity, continuum transport, and statistical thermodynamics. We gathered quantitative experimental data to address the questions above and developed a theoretical model – introducing a stochastic thermal fluctuation term in the energetics of membrane poration – which recapitulates the essential qualitative features of the experimental observations, emphasizes the importance of dynamics, and places the heretofore neglected contribution of thermal fluctuations in driving osmotic response of stressed vesicular compartments.

## Results

### Homogeneous GUVs display swell-burst cycles in hypotonic conditions

The experimental configuration is similar to that already described (Angelova et al., 1992; Oglecka et al., 2014). Briefly, we prepared GUVs consisting essentially of a single amphiphile, namely 1-palmitoyl-2-oleoyl-sn-1-glycero-3-phosphocholine (POPC), doped with a non-perturbative small concentration (1 mol%) of a fluorescently labeled phospholipid (1,2-dipalmitoyl -sn-glycero-3-phosphoethanolamine-N-(lissamine rhodamine B sulfonyl) or Rho-DPPE using standard electroformation technique (Angelova et al., 1992). The obtained GUVs, (typically between 7 and 20 *μ*m in diameter) encapsulated 200 mM sucrose and were suspended in the isotonic glucose solution of identical osmolarity. Diluting the extra-vesicular dispersion medium with deionized water produces a hypotonic bath depleted in osmolytes, subjecting the GUVs to osmotic stress. Shortly (∼1 min delay) after subjecting the GUVs to the osmotic differential, GUVs were monitored using time-lapse epifluorescence microscopy at a rate of 1 image per 150 ms, and images were analyzed using a customized MATLAB code to extract the evolution of the GUVs radius with time, with a precision of about 0.1 *μ*m. A selection of snapshots, revealing different morphological states, and a detailed trace showing the time-dependent changes of vesicle radius and corresponding area strain are shown in Fig. 1b, c, and d, for a representative GUV. Swelling phases are characterized by a quasi-linear increase of the GUV radius (*R*), while pore openings cause a sudden decrease of the vesicle radius.

**Figure 1:**
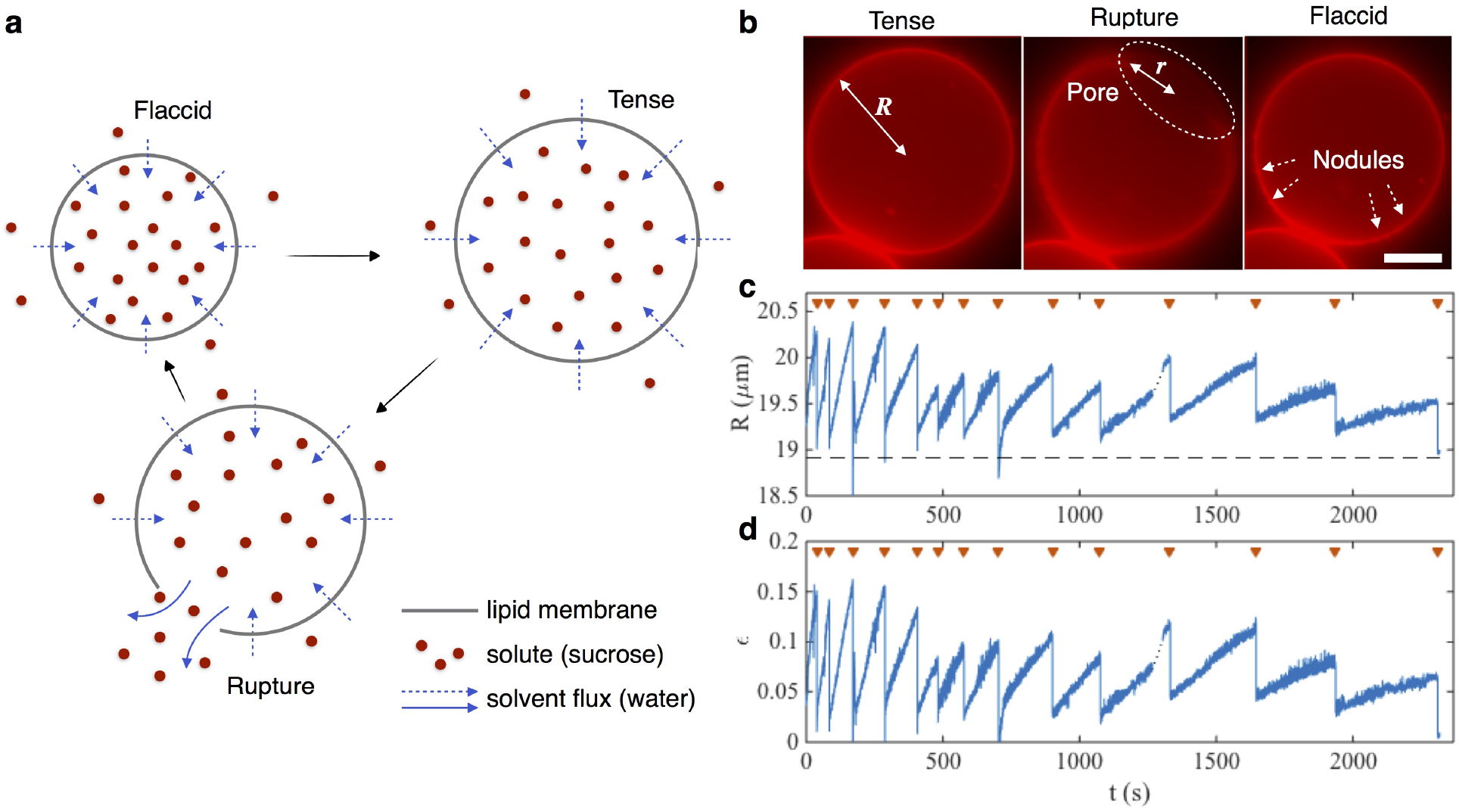
Homogeneous giant unilamellar vesicles (GUVs) made of POPC with 1 mol % Rho-DPPE exhibit swell-burst cycles when subject to hypotonic conditions. **a**, Schematic of a swell-burst cycle of a homogeneous GUV under hypotonic conditions. Blue arrows represent the leak-out of the inner solution through the transient pore. **b**, Micrographs of a swelling (left), ruptured (middle) and resealed (right) GUVs. Scale bar represents 10 *μ*m. Pictures extracted from Supplementary Movie 1. **c**, Typical evolution of a GUV radius with time during swell-burst cycles in 200 mM sucrose hypotonic conditions. The GUV radius increases continuously during swelling phases, and drops abruptly when bursting events occur. Pore opening events are indicated by 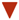. Dash line represents the estimated initial radius *R*_0_. See also Supplementary Movie 2. More GUV radius measurements are shown in Supplementary Fig. 1.d, Computed area strain (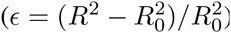).

We outline here three key observations about the dynamics of swell-burst cycles from these experiments.

1. The period between two consecutive bursting events increases with each cycle, starting from a few tenths of a second for the early cycles, to several hundreds of seconds after the tenth cycle.
2. The maximum radius and therefore the maximum strain at which a pore opens decreases with cycle number, suggesting that lytic tension is a dynamic property of the membrane.
3. The observed transient pores are short lived, stay open for about hundred milliseconds, and reach a maximum radius of up to 60 % of the GUV radius.

We seek to explain these observations through a quantitative understanding of the pulsatile GUVs in hypotonic conditions. To do so, we first investigate the mechanics of pore nucleation and its relationship to the GUV swell-burst dynamics.

### Thermal fluctuations drive the dynamics of pore nucleation

In the framework of classical nucleation theory, the energy potential *V* (*r*, *ε*) of a pore in a tense lipid membrane is the balance of two competitive terms, the stretch energy – induced by the membrane tension *σ* – tends to favor the opening and enlargement of a pore, and the edge energy *γ* – originating from the exposure of the hydrophobic lipid tails to the aqueous solution – tends to close the pore (Litster, 1975). The opposing effects of these two quantities produces an energy barrier that the system has to overcome in order for a pore to nucleate. The energy required to open a circular pore of radius *r* in a tensed GUV is represented in Fig. 2a, and b, where the energy barrier is located at a critical pore radius of *r*_*b*_=*γ*/*σ*. When a linear relationship between the stress and the strain is assumed, the height of the energy barrier and the corresponding pore radius are dependent on the membrane strain; the more the membrane is stretched, the lower the energy barrier is, and the smaller the amount of energy required to nucleate a pore.

**Figure 2:**
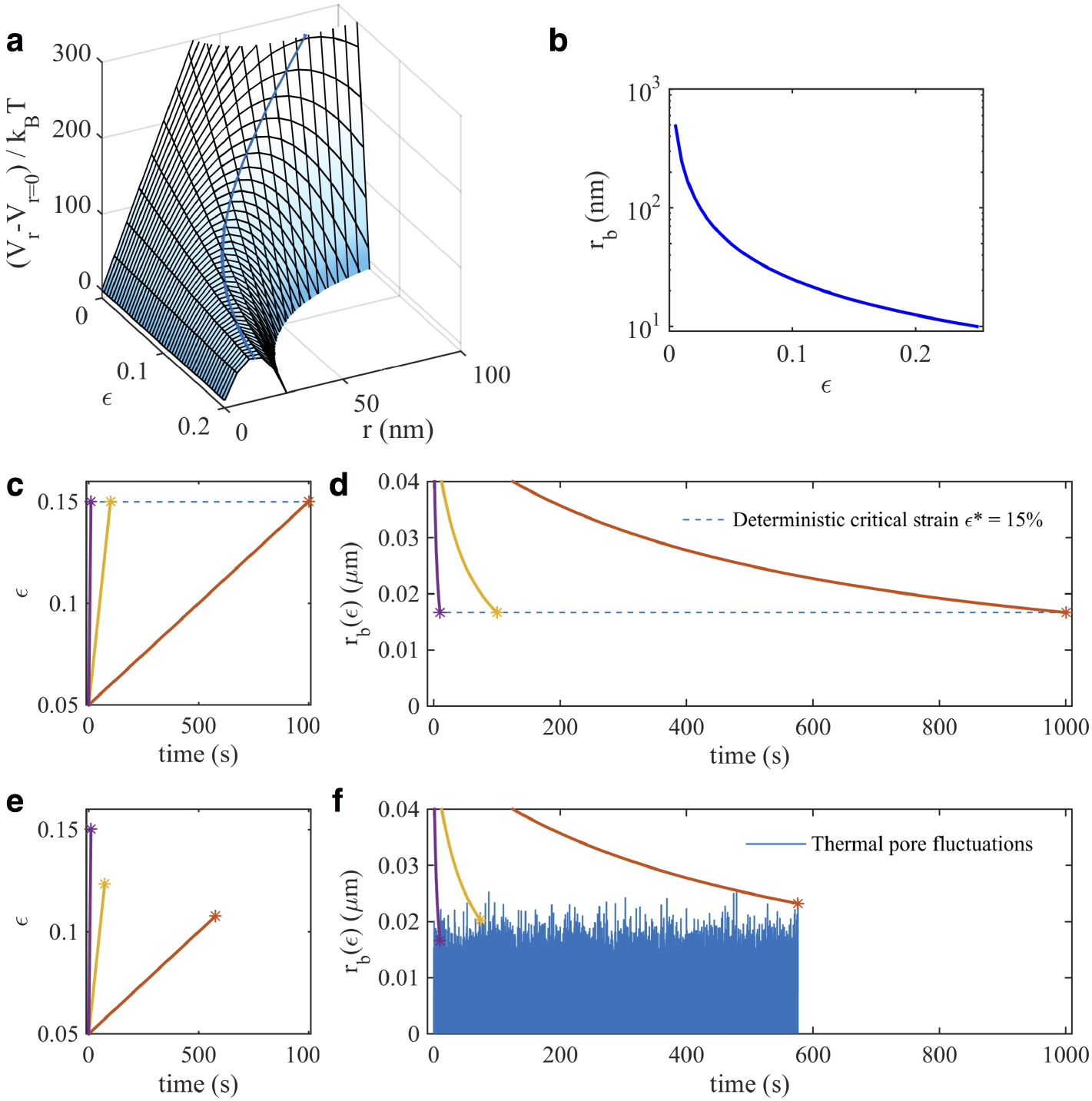
Lytic tension is a dynamic quantity governed by thermodynamic fluctuations. **a**, The energy required to open a pore of radius *r* in a GUV without fluctuations, for various membrane tensions. The energetic cost to open a pore in a tense GUV shows a local maximum, which has to be overcome in order for a pore to open. **b**, For a given strain *ε*, the energy barrier is located at a pore radius *r*_*b*_=*γ*/*σ* (*ε*). **c, d,** Prescribing a linear strain rate (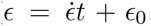, with 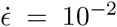, 10^−3^, and 10^−4^ s^−1^, *ε* _0_=0.05), in the deterministic approach, a pore is nucleated whenever the strain reaches a defined lytic strain (c) corresponding to a constant *r*_*b*_ (d), and therefore producing a constant lytic tension. **e, f,** In the stochastic approach however, the nucleation threshold is replaced by a fluctuating pore, inducing a dependence of the lytic strain on the strain rate (e). This is due to the fact that, for lower strain rates, the probability of a large pore fluctuation to reach *r*_*b*_ is higher (f), producing a lower lytic tension on average.

The amplitude of this barrier is strictly positive for finite stretch values, making nucleation impossible without the addition of external energy. This issue has been often resolved by assuming a predetermined and *constant* lytic membrane tension, corresponding to a critical energy barrier under which the pore opens (Fig. 2c, d). However, this approach is in contradiction with our experimental observations that the lytic tension – proportional to the strain in the membrane – varies with each swell-burst cycle (Fig. 1d), due to a dependence on the strain rate (Evans et al., 2003). In order to account for this variation, we include thermal pore fluctuations associated with the nucleation barrier in our analysis. In this scenario, increasing the membrane tension of the vesicle reduces the minimum pore radius *r*_*b*_ at which a pore opens (Fig. 2a, b), lowering the energy barrier down to the range of thermal fluctuations, eventually letting the free energy of the system to overcome the nucleation barrier (Fig. 2e, f). The stochastic nature of the fluctuations can then explain a distribution of pore opening tensions, eliminating the need to assume constant lytic tension.

A direct consequence of the fluctuation activated pore nucleation is that the membrane rupture properties become dynamic. Indeed, fluctuations naturally cause the strain to rupture to be dependent on the *strain rate*, as illustrated in Fig. 2e. In order to understand this dynamic nucleation process, consider stretching the membrane at different strain rates 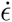. Doing so decreases the radius of the nucleation barrier at corresponding speeds, as shown in Fig. 2f. For slow strain rates, as *r*_*b*_ tends to zero, it spends more time in the accessible range of the thermal pore fluctuations, increasing the probability of a fluctuation to overcome the energy barrier. On the other hand, at faster strain rates, *r*_*b*_ decreases quickly, reaching small values in less time, lowering the probability for above average fluctuation to occur during this shorter time.

We use a Langevin equation to capture the stochastic nature of pore nucleation and the subsequent pore dynamics. This equation includes membrane viscous dissipation, a conservative force arising from the membrane potential, friction with water, and thermal fluctuations for pore nucleation (see Material & Methods for detailed derivation). This yields the stochastic differential equation for the pore radius *r*

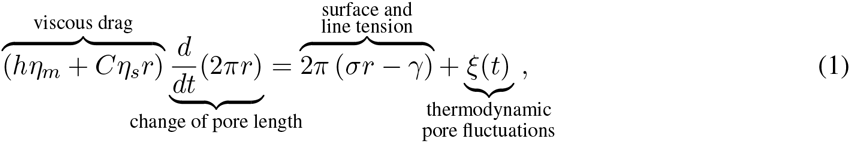

where the noise source *ξ*(*t*) has zero mean and satisfies, 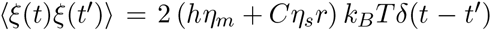 according to the fluctuation dissipation theorem (Kubo, 1966). Here, *η*_*m*_ and *η*_*s*_ are the membrane and solute viscosities respectively, *h* is the membrane thickness, *C* is a geometric coefficient (Ryham et al., 2011; Aubin and Ryham, 2016), *k*_*B*_ is the Boltzmann constant and *T* is the temperature. The values of the various parameters are given in Supplementary Table 1.

### Model captures experimentally observed pulsatile GUV behavior

In addition to pore dynamics (Eq. (1)), we need to consider mass conservation of the solute and the solvent. We assume that the GUV remains spherical at all times and neglect spatial effects. The GUV volume changes because of osmotic influx through the semi-permeable membrane and the leak-out of the solvent through the pore. The osmotic influx is the result of two competitive pressures, the osmotic pressure driven by the solute differential 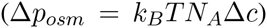, and the Laplace pressure, arising from the membrane tension 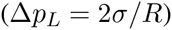, resulting in the following equation for the GUV radius *R*:

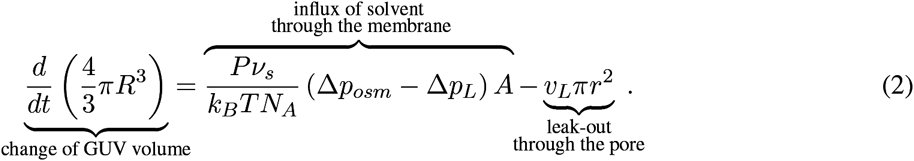

Here *A*=4*πR*^2^ is the membrane area, *P* is the membrane permeability to the solvent, *ν*_*s*_ is the solvent molar volume, and *N*_*A*_ is the Avogadro number. Assuming low Reynolds number regime, the leak-out velocity is given by *v*_*L*_=Δ*pL*^*r*^/(3*πη*_*s*_) (Happel and Brenner, 1983; Aubin and Ryham, 2016).

Mass conservation of solute in the GUV is governed by the diffusion of sucrose and convection of the solution through the pore, which gives the governing equation for the solute concentration differential Δ*c*:

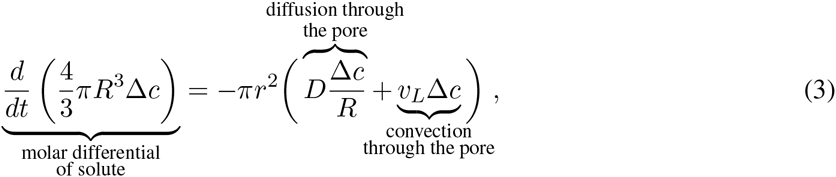

where *D* is the solute diffusion coefficient. These three coupled equations (1)-(3) constitute our model.

In order to completely define the system, we need to specify the relationship between the membrane surface tension *σ* and the area strain of the GUV. We note that the GUV has irregular contours during the pore opening event and for a short time afterwards, when “nodules” are observed at the opposite end from the pore, indicating accumulation of excess membrane generated by pore formation (Fig. 1b middle and right panels). In the low tension regime, GUVs swell by unfolding these membrane nodules, and the stretching is controlled by the membrane bending modulus *κ*_*b*_ and thermal energy, yielding an effective “unfolding modulus” 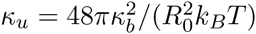 of the order of 10^−5^ N/m (Brochard et al., 1976). In contrast, in the high tension regime, elastic stretching is dominant, and the elastic area expansion modulus *κ*_*e*_ is roughly equal to 0.2 N/m. Since the maximum area strain plotted in Fig. 1d, is about 15 %, significantly larger than the expected 5 % for a purely elastic membrane deformation, the experimental data suggests the occurrence of two stretching regimes: an unfolding driven stretching, and an elasticity driven stretching (Ertel et al., 1993; Hallett et al., 1993; Karatekin et al., 2003). Therefore, for simplicity, we propose a linear dependence between the membrane tension and the strain (*σ* = *κ*_eff_*ε*), where the effective stretching modulus *κ*_eff_ is the only adjustable parameter of the model.

We solved the three coupled equations (1)-(3) for an initial inner solute concentration of *c*_0_ = 200 mM, and different GUV radii of *R*_0_ = 8, 14 and 20 *μ*m. All the results presented here are obtained for *κ*_eff_ = 2 × 10^−3^N/m, the value that best fits the experimental observations (see Supplemental Fig. 5 for the effect of this parameter on the GUV dynamics). Dynamics of the GUV radius and the pore radius are shown in Fig. 3 for a typical simulation with *R*_0_ = 14 *μ*m (see Supplemental Fig. 3 for simulations with different values of *R*_0_). Our model qualitatively reproduces the dynamics of the GUV radius during the swell burst cycle (compare Figs. 1c and 3a). Importantly, we recover the key features of the swell-burst cycle – namely an increase of the cycle period with each bursting event (point 1), and a decrease of the maximum radius with time (point 2). The stochastic nature of the thermodynamic fluctuations leads to variations and occasional irregularities in the pore opening events, and therefore, the cycle period and maximum strain. The dynamics of a single cycle is shown in Fig. 3b,d. Our numerical results show a abrupt drop in the GUV radius, followed by a slower decrease, suggesting a sequence of two leak-out regimes: a fast-burst releasing most of the membrane tension, and a low tension leak-out. This two-step tension release is confirmed by the pore radius dynamics, which after suddenly opening (release of membrane tension), reseals quasi-linearly due to dominance of line tension compared to membrane tension in Eq. (1). Furthermore, the computed pore amplitude and lifetime are in agreement with experimental observations (point 3). Overall, our model is able to reproduce the quantitative features of GUV response to hypotonic stress.

**Figure 3:**
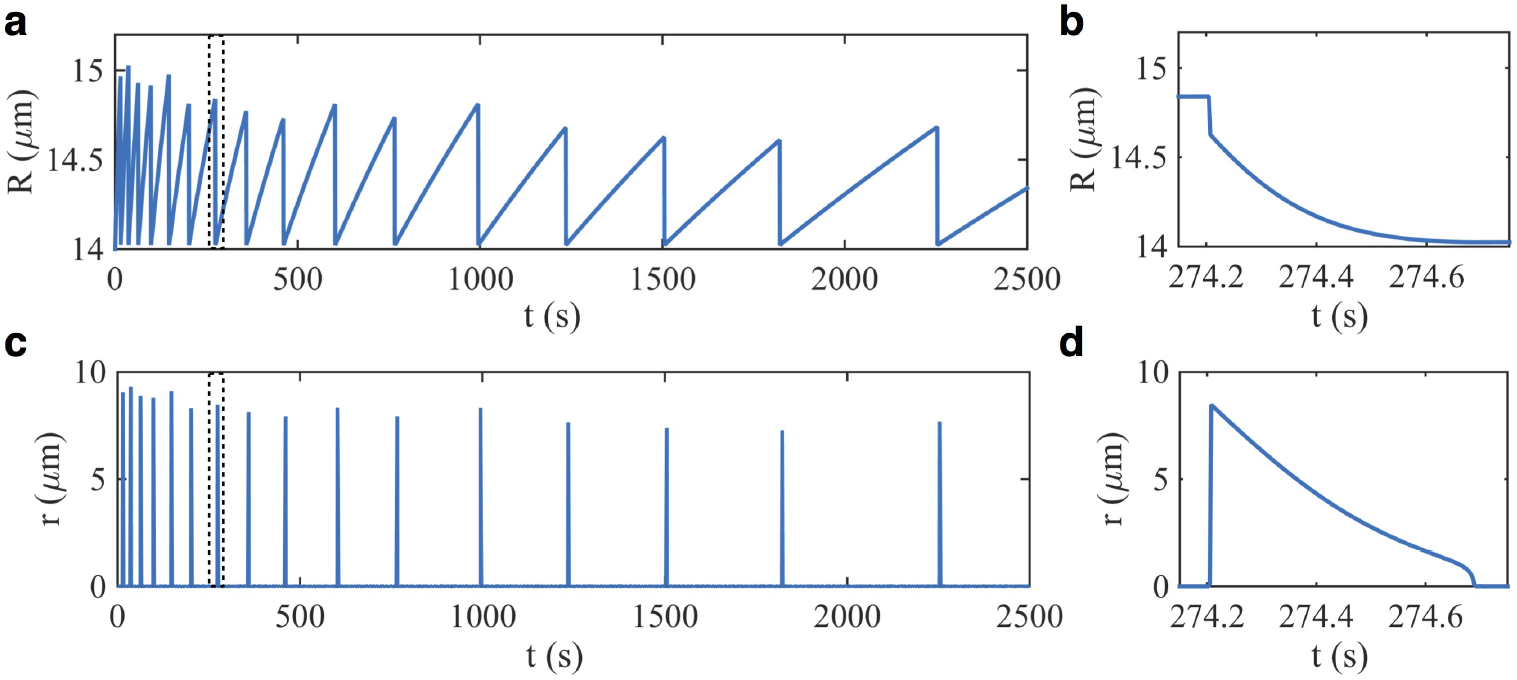
Dynamics of swell-burst cycles from the model for a GUV of radius 14 *μ*m in 200 mM hypotonic stress. **a, b,** GUV radius and **c, d,** pore radius as a function of time. **a, c,** The model captures the dynamics of multiple swell-burst cycles, in particular the decrease of maximum GUV radius and increase of cycle period with cycle number. **b, d,** Looking closely at one pore opening event corresponding to the region between the dashed lines, the model predicts a three stage pore dynamics (opening, closing, resealing), with a characteristic time of few hundreds milliseconds. Numerical reconstruction of the GUV is shown in Supplementary Movie 3 and 4. Results for *R*_0_ = 8 and 20 *μ*m are shown in Supplementary Fig. 3.

If thermal fluctuations are ignored, the strain to rupture needs to be adjusted to roughly 15% in order to match the range of maximum GUV radius observed experimentally (Supplementary Fig. 4). However such a deterministic model does not capture the pulsatile dynamics as well as the stochastic model, failing to reproduce a strain rate dependent maximum stress (Supplementary Fig. 11).

### Solute diffusion is dominant during the low tension regime of pore resealing

The concentration differential of sucrose decreases exponentially and drops from 200 mM to about 10 mM in about 1000 seconds (Supplementary Fig. 3). Even after 2000 s when the concentration differential is as low as 10 mM, the osmotic influx is still large enough to maintain the dynamics of swell-bursts (Fig. 1c, Supplementary Fig. 3). We further observe that every pore opening event produces a sudden drop in inner concentration (Fig. 4a, blue line). This suggests that diffusion of sucrose plays an important role in governing the dynamics of solute. In the absence of diffusive effects, the model does not show the abrupt drops in concentration but a rather smooth exponential decay (Fig. 4a, grey line).

**Figure 4:**
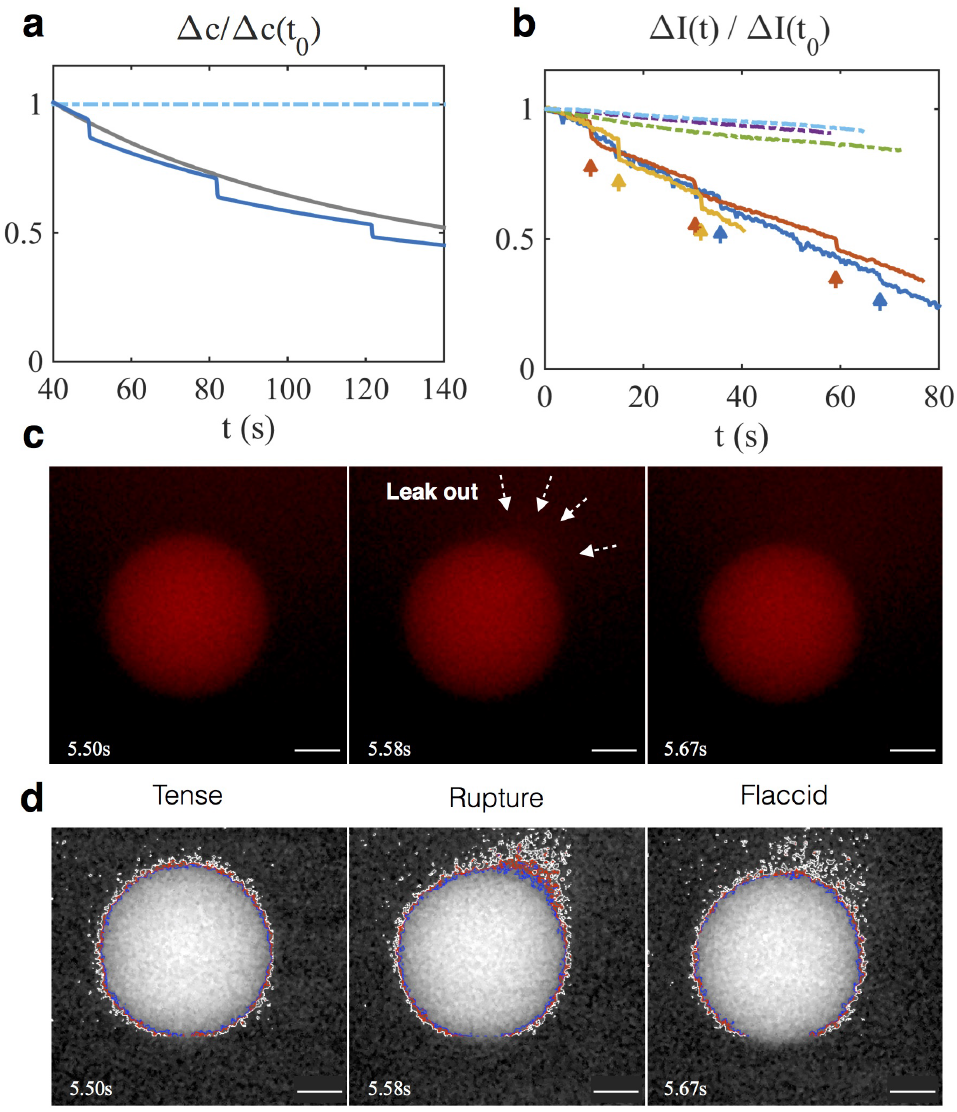
Solute dynamics. **a,** Normalized concentration differential (here *t*_0_ = 40 s) with diffusion (blue line), without diffusion (grey line), and in the absence of osmotic differential (dashed line). The concentration drops rapidly at each bursting event due to diffusion, while in the absence of diffusion, there are no concentration drops over time and the profile is smooth (see Supplementary Fig. 6 for further analysis on the effect of diffusion). **b,** Time evolution of the normalized fluorescence intensity of a GUV in hypotonic condition, encapsulating fluorescent glucose analog. Δ*I* is the difference in mean intensity between the inside of the GUV and the background. When in hypotonic conditions (solid lines) the normalized intensity decreases with time due to the constant influx of water through the membrane, and shows sudden drops in intensity at each pore opening (indicated by arrows), due to diffusion of sucrose through the pore (see Supplementary Movie 5). In comparison, GUVs in isotonic condition (dashed lines) exhibit a rather constant fluorescence intensity (see Supplementary Movie 6). **c,** Micrographs of a GUV in hypotonic condition, encapsulating fluorescent glucose analog, just prior bursting (left panel), with an open pore (middle panel), and just after pore resealing (right panel). The leak-out of fluorescent dye is observed in the middle frame, coinciding with a drop of the GUV radius. Frames extracted from Supplementary Movie 7. **d,** Same as panel (c), with the images processed to increase contrast and attenuate noise. The blue, red, and white lines are the isocontours of the 90, 75 and 60 grey scale values respectively, highlighting the leak-out of fluorescent dye.

To experimentally verify the model predictions of sucrose dynamics, we quantified the evolution of fluorescence intensity in GUVs encapsulating 200 mM sucrose plus 58.4 *μ*M 2-NBDG, a fluorescent glucose analog (see Material and Methods). Fig. 4b presents the evolution of fluorescent intensity of sucrose in time. GUVs in isotonic conditions (dashed lines) do not show a significant change in fluorescence intensity. GUVs in hypotonic conditions (solid lines) exhibit an overall decrease of intensity due to permeation of water through the membrane. Strikingly, consecutive drops of fluorescence intensity are observed coinciding with the pore opening events (Fig. 4c,d middle panels), and point out the importance of sucrose diffusion through the pore. While the quantitative dynamics of sucrose depends on the value of the diffusion constant (Supplementary Fig. 6), the qualitative effect of diffusion on the dynamics remains unchanged. On the other hand, leak-out induced convection does not influence the inner concentration of sucrose, as both solvent and solute are convected, conserving their relative amount. These observations are in agreement with the existence of the low tension pore closure regime discussed above, where Laplace pressure produces negligible convective transport compared to solute diffusion though the pore.

### Cycle period and strain rate are explicit functions of the cycle number and GUV properties

Given that lytic tension is a dynamic quantity, we asked how cycle period and strain rate evolve along the cycles. We studied the dynamics of GUVs with resting radii of 8, 14 and 20*μm*, each simulation data point representing the mean and the standard deviation (because of the stochastic nature of the model, 10 simulations per GUV radius were run). Cycle periods and strain rates show a dependence on the GUV radius, as depicted in Fig. 5a-d. Indeed, larger GUV have slower dynamics, resulting in smaller strain rates and longer cycle period (Supplementary Fig. 3). To verify this experimentally, a total of eight GUVs were analyzed with resting radii ranging from 7.02 to 18.76 *μ*m (Supplementary Fig. 1). The measured cycle period and strain rate as a function of the cycle number (corrected for the lag between the application of the hypotonic stress and the beginning of the observations) are showed in Fig. 5a, and b, respectively. Experimental and model results quantitatively agree, and show a exponential dependence of the cycle period and strain rate on cycle number (Insets Fig. 5a,b).

**Figure 5:**
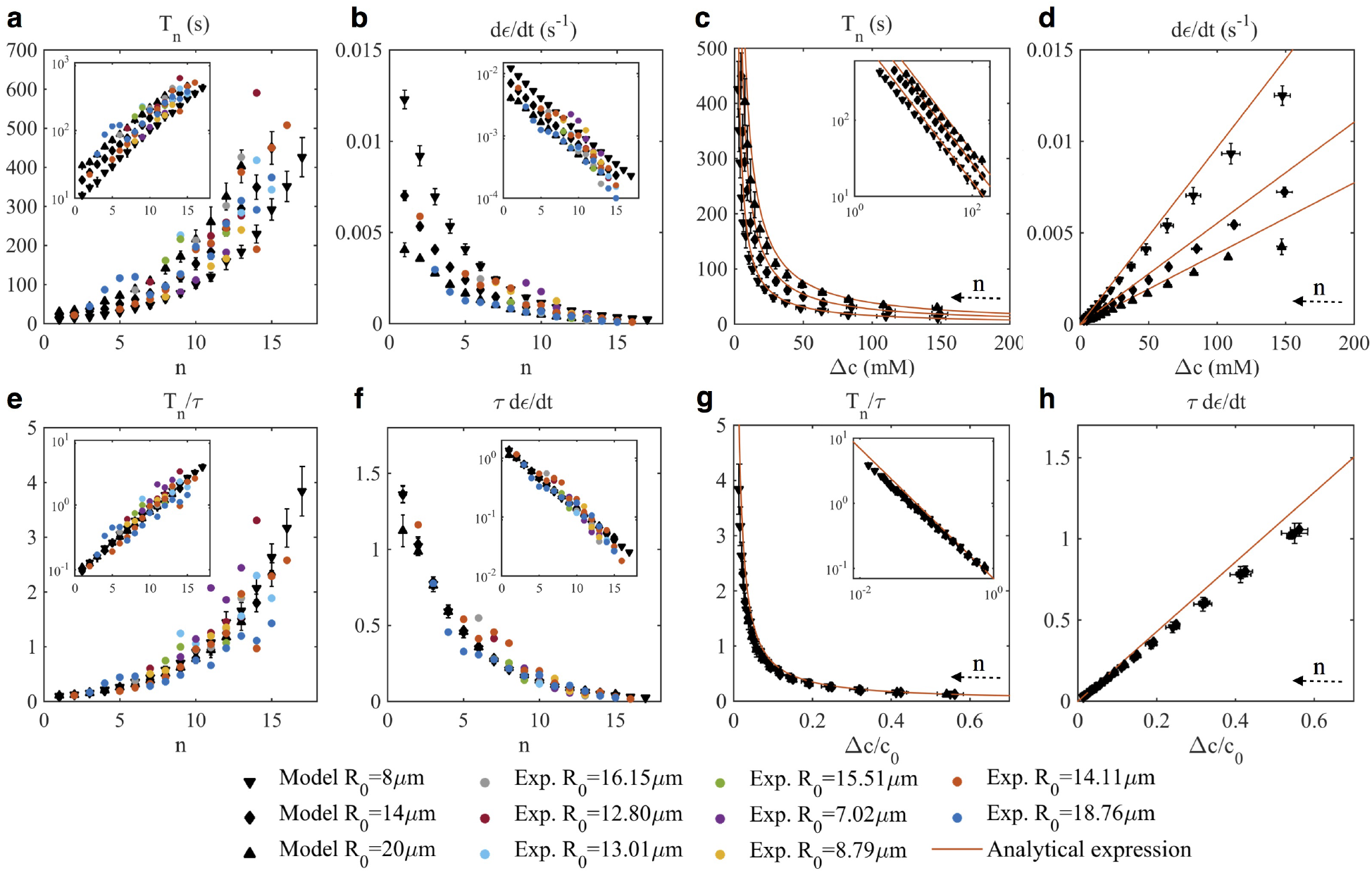
Cycle period and strain rate are exponential function of cycle number, and power-law functions of solute concentration. **a, c, e, g,** Cycle period and **b, d, f, h,** strain rates as functions of cycle number (*n*) **a, b, e, f,** and solute concentration **c, d, g, h. e-h,** are data from **a-d,** scaled by the characteristic time associated with swelling defined as 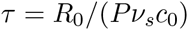. Insets show the same data in log scale. Each model point is the mean of 10 numerical experiments, error bars represent±standard deviations. **e-h,** The nondimensionalization by τ allows cycle periods and the strain rates to collapse onto a single curves. The analytical expressions for the cycle period 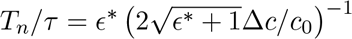, and strain rate 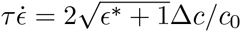 with 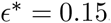, are plotted for comparison (see Supplementary Material for their derivation).

Two further questions arise: How can we relate the cycle number to the driving force of the process, that is to say the osmotic differential? And, is there a scaling law that governs the GUV dynamics? Hence we computed the cycle solute concentration (defined as the solute concentration at the beginning of each cycle) as a function of the cycle number (Supplementary Fig. 2). We found that the solute concentration follows an exponential decay function of the cycle number, and is independent of the GUV radius. Additionally, plotting the cycle period and strain rate against the cycle solute concentration (Fig. 5c,d), we observe that the cycle period decreases as the inverse of the concentration, while the strain rate is a linear function of Δ*c*. The data presented in Fig. 5a-d suggest that the dynamics of GUVs swell-burst cycle can be scaled in some way to their size. From the non-dimensional form of Eq. (2), we extracted a characteristic time associated with swelling, defined by 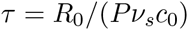, and scaled the cycle period and strain rates with this quantity. As shown in Fig. 5e-h, all the scaled experimental and model data collapse onto the same curve, within the range of the standard deviations. The scaled relationships can be justified analytically, by estimating the strain rate and cycle period as 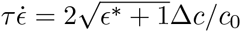, and 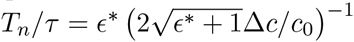 respectively (see Supplementary Material for full derivation). These analytical expressions are plotted in Fig. 5c, d, g, h for a characteristic critical strain of 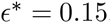, showing good agreement with the numerical data. Taken together, these results suggest that the GUV pulsatile dynamics is governed by the radius, the membrane permeability, the solute concentration, and importantly the pore nucleation mechanism which determines the strain to rupture.

## Discussion

Explaining how membrane-enclosed compartments regulate osmotic stress is a first step towards understanding how cells control volume homeostasis in response to environmental stressors. In this work, we have used a combination of theory and experiments in a simple model system to study how swell-burst cycles control the dynamics of GUV response to osmotic stress. Using this system, we show that the pulsatile dynamics of GUVs under osmotic stress is controlled through thermal fluctuations that govern pore nucleation and lytic tension.

The study of osmotic response of lipid vesicles has a rich history in biophysics, starting with the pioneer theoretical work by Koslov and Markin (1984). The central feature of GUV osmotic response is the nucleation of a pore. Yet, while Evans and coworkers identified that rupture tension was not governed by an intrinsic critical stress, but rather by the load rate, the idea of a constant lytic tension has persisted in the literature (Idiart and Levin, 2004; Popescu and Popescu, 2008; Peterlin and Arrigler, 2008). By coupling fluctuations to pore energy, we have now reconciled the dynamics of the GUV over several swell-burst cycles with pore nucleation and dependence on strain rate. Our model is not only able to capture the experimentally observed pulsatile dynamics of GUV radius and solute concentration (Figs. 3 and 4), but also predicts pore formation events and pore dynamics (Fig. 3c,d). We also found that during the pore opening event, a low-tension regime enables a diffusion dominated transport of solute through the pore (Fig. 4), a feature that has been up to now neglected in the literature.

Specifically, we have identified a scaling relationship between (i) the cycle period and cycle number and (ii) the strain rate and the cycle number, highlighting that swell-burst cycles of the GUVs in response to hypotonic stress is a dynamic response (Fig. 5). One of the key features of the model is that we relate the cycle number, an experimentally observable quantity, to the concentration difference of the solute, a quantity that is hard to measure in experiments (Supplementary Fig. 2). This allows to interpret the scaling relationships described above in terms of solute concentration differential. The cycle period shows an inverse relationship with the solute concentration, while the strain rate is a linear function of the concentration difference. Both relationships are derived theoretically in the Supplemental material. These features indicate long time scale relationships of pulsatile vesicles in osmotic stress.

Thermal fluctuations and stochasticity are known to play diverse roles in cell biology. Now well recognized examples include Brownian motors and pumps (Jülicher et al., 1997; Oster, 2002), noisy gene expression (Elowitz et al., 2002), and red blood cell flickering (Turlier et al., 2016). The pulsatile vesicles system presented here provides yet another example of how fluctuations can be utilized by simple systems to produce dynamical adaptive behavior. Given the universality of fluctuations in biological processes, it appears entirely reasonable that simple mechanisms similar to the pulsatile vesicle have been exploited by early cells, conferring them with a thermodynamic advantage against the environmental osmotic assaults.

While we have been able to explain many fundamental features of the pulsatile GUVs in response to osmotic stress, there are some obvious limitations of our approach and need for further experiments. We have assumed a linear relationship between stress and strain. Although this assumption is reasonable and appears to work well for the present experimental conditions, a more general expression should be considered to include both membrane (un)folding and elastic deformation (Helfrich and Servuss, 1984). Another important aspect of biological relevance is membrane composition, where the abundance of proteins and heterogeneous composition leading to in-plane ordering and asymmetry across leaflets influence the membrane mechanics (Alberts et al., 2014; Rangamani et al., 2014). We have previously found experimentally that the dynamics of swell-burst cycles is related to the compositional degrees of freedom of the membrane (Oglecka et al., 2014). Future efforts will be oriented toward the development of theoretical framework and quantitative experimental data that provide insight into how membrane’s compositional degrees of freedom influences membrane mechanics.

## Acknowledgments

We are grateful to Prof. Wouter-Jan Rappel and Prof. Alex Mogilner for insightful comments on the manuscript. We also thank Prof. Daniel Tartakovsky for enriching discussions. This work was supported in part by the FISP 3030 for the year 2015-2016 to M.C., NTU provost office to J.C.S.H., AFOSR FA9550-15-1-0124 award to P.R., and NSF PHY-1505017 award to P.R. and A.N.P.

## Material and Methods

### Experimental Material and Methods

#### Swell-burst cycle experiments

The experimental methods for the GUVs preparation has been described in (Oglecka et al., 2014; Angelova et al., 1992). Briefly, GUVs (100% POPC+1mol% Rho-DPPE) containing 200 mM of sucrose were prepared by electroformation, yielding vesicles with radii ranging from 7 to 20 *μ*m. GUVs were then placed in a bath of deionized water at room temperature, inducing hypotonic stress proportional to the inner sucrose concentration. The kinetics of eight GUVs were recorded by time-lapse microscopy at 1/150 images/ms. In order to allow for the sedimentation of GUVs to the bottom of the well, observations were started about one minute after the GUVs were subject to hypotonic conditions.

For each frame, the GUV radius was measured using a customized MATLAB (Mathworks, Natick, MA) code to streamline the image analysis. This code uses a Circular Hough Transform method based on a phase-coding algorithm to detect circles (Davies, 2012), and measure their radii and centers. For our data, this custom code gives the evolution of the GUV radius in time with a precision of about 0.1 *μ*m. Due to slow movement of the GUVs, in some cases the observation fields had to be adjusted to follow the GUVs, and the recording was paused. These are indicated by black dashed lines in Supplementary Fig. 1. In order to define a systematic experimental initial GUV radius, R0 was determined for each GUV as 0.995 times the first measured local minimum GUV radius, in accordance with our numerical results. Furthermore, burst events were identified by drops of GUV radius larger than 0.2 _m within a 1.5 s interval, and are plotted as solid red triangle. Bursting events that were likely to happen during the video gaps were indicated by plain red triangle (these “likely” bursting events were not taken into account in the data processing for Fig. 5).

#### Leak-out quantification

To quantify the leak-out amount when a membrane pore is formed, giant unilamellar vesicles (GUVs) were electroformed in 200 mM sucrose, supplemented with 58.4 *μ*M 2-NBDG (2-(N- (7-Nitrobenz-2-oxa-1,3-diazol-4-yl) Amino) -2-Deoxyglucose), a fluorescent glucose analog that has an almost identical molecular weight as sucrose. Fluorescence imaging was performed on a deconvolution microscope, equipped with a FITC filter. Time-lapse imaging of the vesicles was performed approximately one minute after exposing the vesicles to either deionized water (hypotonic conditions *n* = 3) or glucose (isotonic conditions *n* = 3) environment to ensure sedimentation of GUVs to the bottom of the well. All acquisitions were performed using identical settings to facilitate comparison of vesicles submerged in water or equi-osmotic glucose environment.

For Fig. 4c, the GUVs were detected with a MATLAB (Mathworks, Natick, MA) code adapted from the one described above, where the mean gray intensity inside and outside of the GUV are measured. For every time frame, the difference between the inner and outer mean intensity Δ*I*(*t*) was computed, and normalized by the intensity difference of the first frame Δ*I*(*t*_0_). Bursting events were identified by visual inspection of the videos, and reported by arrows on Fig. 4b.

In order to highlight the efflux of fluorescent dyes during a GUV bursting event, three frames (before, during and after the event) were extracted form the video of a GUV containing 200 mM sucrose+58.4 *μ*M 2-NBDG in hypotonic conditions (Fig. 4c). These images were further processed with ImageJ software (Schneider et al., 2012) to plot Fig. 4d. Briefly, the noise was attenuated by successively applying ImageJ built-in routines (background suppression, contrast enhancing, median filter), and ploting the isovalues of gray at 90, 75 and 60 with the pluggin Coutour Plotter^1^.

#### Burst cycle analysis

A swell-burst cycle was defined between two successive minimum GUV radius that immediately followed a bursting event. Cycle periods were computed as the time between two consecutive minima in vesicular radii, when there was no video gaps (two successive solid triangles in Fig. 1c and Supplementary Fig. 1). The strain rate was computed as the difference between the maximum and minimum radii within these cycles, divided by the time between these two events. Because of the lag between the beginning of the experiments and the beginning of the video recordings, the initial observed cycle number was adjusted between *n* = 1 and *n* = 4, depending on *R*_0_.

### Theoretical development

Here we derive a theoretical model to describe the swell-burst cycle of a GUV under hypotonic conditions. In line with previous work (Koslov and Markin, 1984; Idiart and Levin, 2004; Popescu and Popescu, 2008; Peterlin et al., 2012b), the model has three conservation equations, governing the dynamics of the solvent, the solute, and the membrane pore.

#### Mass conservation of solvent

Mass conservation of the solvent (water) within the vesicle is governed by the flux through the membrane (*j*_*w*_), and the leak-out through the pore. For a spherical GUV, the general form of the mass conservation equation for the solvent is

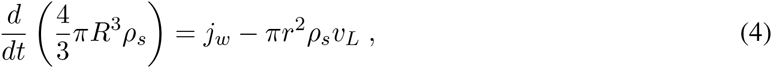

where *R* and *r* are the radius of the vesicle and the pore respectively, *ρ*_*s*_ is the mass density of the solvent, and *v*_*L*_ is the leak-out velocity of the solvent. The osmotic flux is influenced by the permeability of the membrane to the solvent (*P*), the osmotic pressure (Δ_*posm*_), and the Laplace pressure (Δ_*pL*_). A phenomenological expression for the osmotic flux is (Koslov and Markin, 1984; Popescu and Popescu, 2008; Peterlin et al., 2012a)

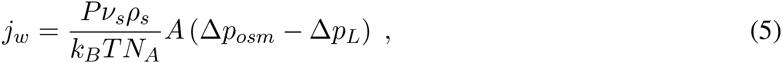

where *ν*_*s*_ is the solvent molar volume, and the membrane area is defined as *A* = 4*πR*^2^−*πr*^2^. The two pressures involved in Eq. (5) are defined as

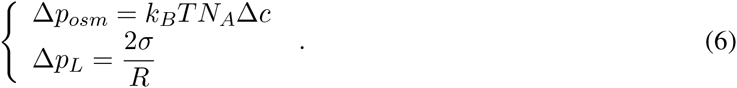

The Laplace pressure originates from the surface tension in the membrane *σ*, which we assume to be proportional to the membrane strain

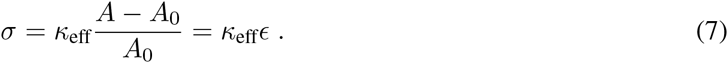

Here *κ*_eff_ is the effective area extension modulus (combining the effects of membrane unfolding and elastic deformation), and *A*_0_ = 4*πR*^2^_0_ is the surface of the vesicle in its unstretched state. The leak-out velocity *v*_*L*_ can be analytically approximated at low Reynolds number in order to relate it to the Laplace pressure (Happel and Brenner, 1983)

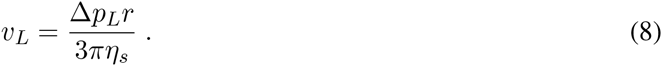

Substituting these definitions into Eq. (4), the mass conservation equation for the solvent takes the form of an ordinary differential equation (ODE) for the GUV radius

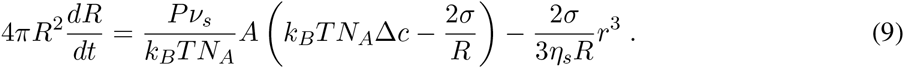

#### Mass conservation of solute

The permeability of lipid membranes to water is several orders of magnitude larger than for most solutes (Fettiplace and Haydon, 1980; Deamer and Bramhall, 1986). Consequently the lipid bilayer is supposed to be semi-permeable, neglecting sucrose transport through the membrane. Thus, variation of solute in the vesicle is exclusively limited to diffusive and convective transport through the pore, such that

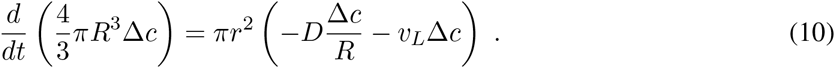

While the diffusive flux through the pore is usually neglected over the convective efflux of solute, theoretical analysis of long lived pores indicates that the Laplace pressure decreases rapidly after the pore opening, and stays low for most of the pore life time (Brochard-Wyart et al., 2000). This suggests that the convective efflux directed by the leak-out velocity may not always be the dominant solute transport mechanism, as confirmed by our numerical and experimental results (see main text Fig. 4). Expanding Eq. (10) we obtain a ODE for the concentration difference in solute

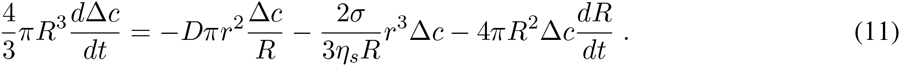

#### Pore force balance

The pore in the lipid bilayer is modeled as an overdamped system, where the pore radius is governed by the following Langevin equation (Seifert, 2008)

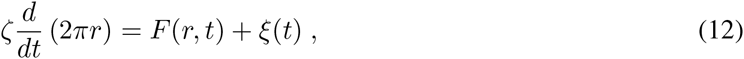

where ζ is the membrane drag coefficient (inverse of the mobility), *F*(*r*; *t*) is a conservative force, and *ξ* is a noise term accounting for independent thermally-induced pore fluctuations. The drag coefficient includes two in-plane contributions ζ = ζ_*m*_ + ζ_*s*_: one from membrane dissipation, proportional to the membrane viscosity and thickness ζ_*m*_ = *η*_*m*_*h* (Brochard-Wyart et al., 2000), and a second from the friction of the solvent with the moving pore – proportional to the solvent viscosity 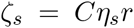, where *C* = 2*π* is a geometric coefficient (Ryham et al., 2011; Aubin and Ryham, 2016). The conservative force 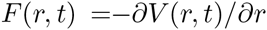 arises from the membrane potential *V* (*r*, *t*), which is equal to the sum of the stretching potential *V*_*s*_, and the pore energy *V*_*p*_. We assume the membrane stretching energy to take a Hookean form 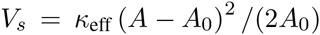, where *κ*_eff_ is an effective stretching modulus approximating the combined contributions of membrane unfolding and elastic stretching. The pore energy depends on the edge energy and length as *V*_*p*_ = 2*πrγ*, where *γ* is the pore line tension, here assumed independent of the pore radius. Using the definition 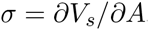, we can therefore express the force as

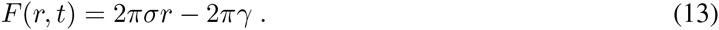

The fluctuation term has a zero mean, and a correlation function given by

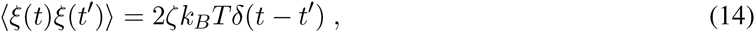

following the dissipation-fluctuation theorem, where *δ* is the Dirac delta function.

Rearranging Eq. (12) with these definitions, we obtain a stochastic differential equation for the pore radius

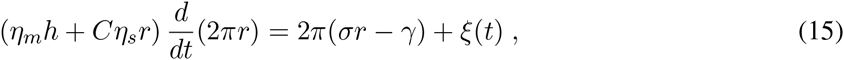

with *r* ≥ 0. The last term in Eq. (15) is responsible for thermally driven pore nucleation.

#### Deterministic model

In the absence of thermal fluctuations, a critical value for the membrane tension (or strain) has to be defined, and an initial pore has to be set artificially in order for a large pore to open. In that case, Eq. (15) is simply

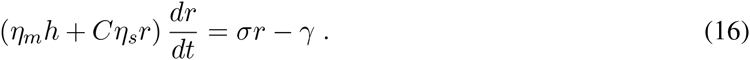

When the pore is closed (*r* = 0) and the strain overcomes the predetermined critical value (*ε* ≥ *ε*\*), an initial pore large enough to overcome the nucleation barrier (*r* = *γ*/*σ*) is artificially created.

### Numerical implementation

All numerical computations have been carried out using custom codes in MATLAB (Mathworks, Natick, MA). The stochastic model, composed by Eqs. (9), (11), and (15) was solved using an order-1 Runge-Kutta scheme. Because a pore nucleation event occurs due to a single fluctuation overcoming the energy barrier, the numerical implementation of the noise requires the definition of a fluctuation frequency *f*_*T*_ that is independent of the time step. The deterministic model (Eqs. (9), (11), and (16)) was solved using Euler method. All parameters are shown in Supplementary Table 1. All time steps were taken as 0.1 ms, (smaller time steps did not improve the accuracy of the results significantly). For the cycles analysis of the stochastic model, Fig. 5, shows the average and standard deviations of 10 runs with same parameters.

## Supplementary Material

### Theroretical derivation of the relation between cycle period, strain rate, and concentration differential

First, we derive the linear dependence of the strain rate on the concentration difference shown in Fig. 5h. For a closed vesicle (*r* = 0), the membrane area is *A* = 4*πR*^2^, and the strain rate is

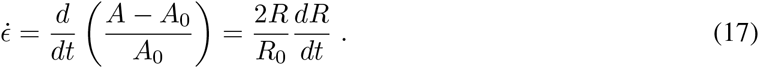

This allows us to write Eq. (9) in terms of the strain rate as

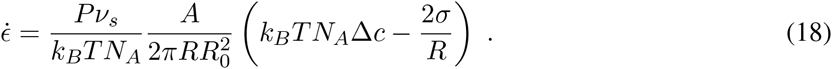

When the osmotic pressure is the dominant process influencing GUV swelling, we can neglect the Laplace pressure and obtain

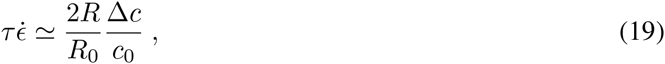

where 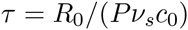. At maximum GUV radius amplitude, *R*/*R*_0_ can be expressed in term of the ctitical strain as 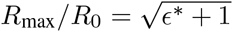, allowing to write Eq. (19) as

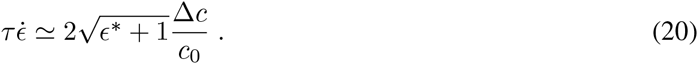

Plotting this relationship in Fig. 5d, h for a typical critical strain *ε*\* = 0.15, we get a good agreement with the numerical results from the stochastic model.

We now derive an approximate relation between the cycle period and the strain rate. During a cycle of period *T*_*n*_, the critical strain can be written

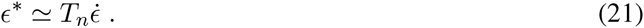

Introducing Eq. (20), we get

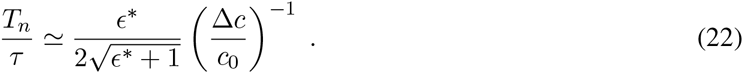

Taking *ε*\* = 0.15, this relationship fits well the simulation results, as shown in Fig. 5c, g.

It should be noted that, because the Laplace pressure is neglected in the derivation of Eq. (20), the analytical expression slightly overestimates the strain rate as shown in Fig. 5d, h. Moreover the cycle period is also overestimated for low solute concentrations due to the constant critical strain assumed in the analytical expression (Fig. 5c, g).

#### Supplementary Movie 1

Membrane nodules appearance after membrane pore reseals. Movie assembled from time-lapse fluorescence microscopy images (frame rate, 2 fps; total duration, 17 s; image size, 82.43 *μ*m × 82.43 *μ*m; scale bar, 10 *μ*m) obtained for a population of electroformed GUVs consisting of POPC doped with 1% Rhodamine-B labeled DPPE membrane in a hypotonic solution (Osmotic differential of 200 mM).

#### Supplementary Movie 2

Multiple swell-burst cycles of GUVs subject to hypotonic stress. Movie assembled from time-lapse fluorescence microscopy images (frame rate, 24 fps; total duration, 77 s; image size, 82.43 *μ*m × 82.43 *μ*m; scale bar, 10 *μ*m) obtained for a population of electroformed GUVs consisting of POPC doped with 1% Rhodamine-B labeled DPPE membrane in a hypotonic solution (Osmotic differential of 200 mM).

#### Supplementary Movie 3

Model results showing multiple swell-burst cycles of a GUV subject to hypotonic stress. GUV radius (top-left panel), pore radius (middle-left panel), and solute differential (bottom-left panel) as a function of time. Right panel is a representation of the numerical GUV in time, where the grey intensity is proportional to the inner sucrose concentration. GUV initial radius is *R*_0_ = 14 *μ*m, initial solute concentration is *c*_0_ = 200 mM. All parameters are shown in Supplementary Table 1.

#### Supplementary Movie 4

Model results showing a single pore opening dynamics of a GUV subject to hypotonic stress. GUV radius (top-left panel), pore radius (middle-left panel), and solute differential (bottomleft panel) as a function of time. Right panel is a representation of the numerical GUV in time, where the grey intensity is proportional to the inner sucrose concentration. GUV initial radius is *R*_0_ = 14 *μ*m, initial solute concentration is *c*_0_ = 200 mM. All parameters are shown in Supplementary Table 1.

#### Supplementary Movie 5

Solute leakage of a GUV in multiple swell-burst cycles under hypotonic condition. Movie assembled from time-lapse fluorescence microscopy images (frame rate, 24 fps; total duration, 11 s; image size, 119.14 *μ*m × 125.58 *μ*m; scale bar, 20 *μ*m) obtained for a population of electroformed GUVs consisting of POPC doped with 1% Rhodamine-B labeled DPPE membrane in a hypotonic solution (Osmotic differential of 200 mM).

#### Supplementary Movie 6

GUV under isotonic condition. Movie assembled from time-lapse fluorescence microscopy images (frame rate, 24 fps; total duration, 8 s; image size, 101.11 *μ*m × 101.11 *μ*m; scale bar, 10 *μ*m) obtained for a population of electroformed GUVs consisting of POPC doped with 1% Rhodamine-B labeled DPPE membrane in a isotonic solution (no osmotic differential).

#### Supplementary Movie 7

Solute efflux from GUV during one swell-burst cycle. Movie assembled from time-lapse fluorescence microscopy images (frame rate, 12 fps; total duration, 8 s; image size, 164.86 *μ*m × 164.86 *μ*m; scale bar, 20 *μ*m) obtained for a population of electroformed GUVs consisting of POPC doped with 1% Rhodamine-B labeled DPPE membrane in a hypotonic solution (Osmotic differential of 200 mM).

**Supplementary Figure 1:**
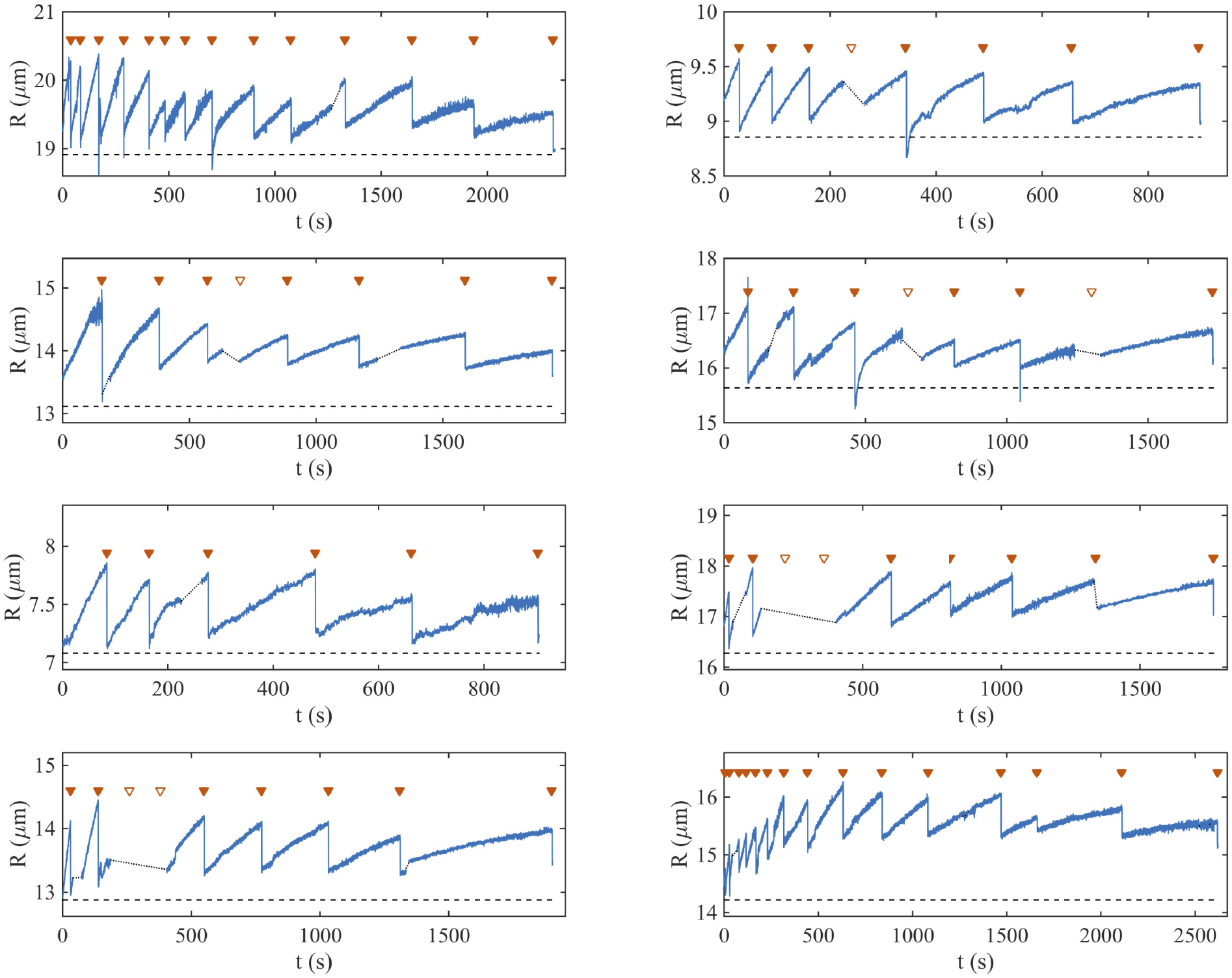
Experimental measurements of GUV radius during swell-burst cycles in 200 mM sucrose hypotonic solutions. Videos of GUV were recorded and analyzed with a contour detection software, as described in the Material and Methods section. The radii of eight GUVs from different experiments are plotted here as a function of time. The radii increase continuously during swelling phases, and drop abruptly when bursting events occur. Each observed pore opening event is indicated by a 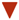. Gaps in the videos due to experimental constraints are shown by dashed lines, and pore opening events that likely occurred during these gaps are indicated by ∇.

**Supplementary Table 1:** Parameters used in the simulations

| Parameter | Typical value | References |
| --- | --- | --- |
| $R_0$ | 8-20 $\mu\text{m}$ | this work |
| $c_0$ | 200 mM | this work |
| $d$ | 3.5 nm | |
| $\rho_s$ | 1000 kg m<sup>-3</sup> | |
| $\nu_s$ | $18.04 \times 10^{-6} \text{ m}^3 \text{ mol}^{-1}$ | |
| $P$ | 20 $\mu\text{m/s}$ | (Olbrich et al., 2000) |
| $T$ | 294 K | |
| $\gamma$ | 5 pN | (Portet and Dimova, 2010) |
| $\kappa_{\text{eff}}$ | $2 \times 10^{-3} \text{ N/m}$ | this work |
| $\eta_m$ | 5 Pa s | (Hormel et al., 2014) |
| $\eta_s$ | 0.001 Pa s | |
| $D$ | $5 \times 10^{-10} \text{ m}^2/\text{s}$ | (Linder et al., 1976) |
| $C$ | $2\pi$ | (Aubin and Ryham, 2016) |
| $f_T$ | 150 Hz | this work |

**Supplementary Figure 2:**
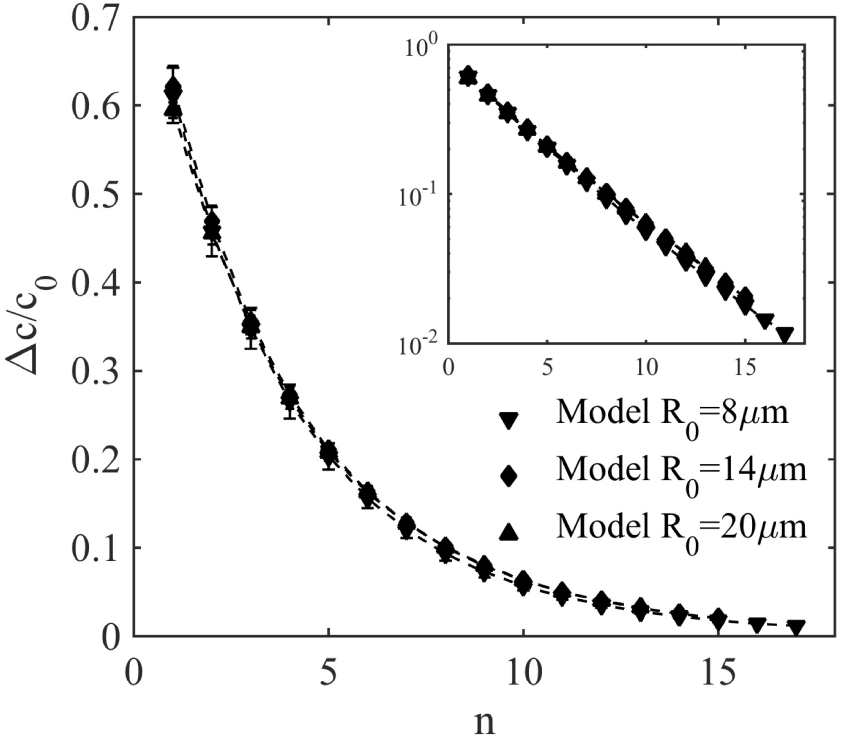
Model results for the solute concentration as a function of the cycle number. Insets shows the same data with a logarithmic y axis, highlighting the exponential dependence 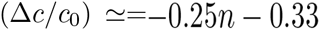.

**Supplementary Figure 3:**
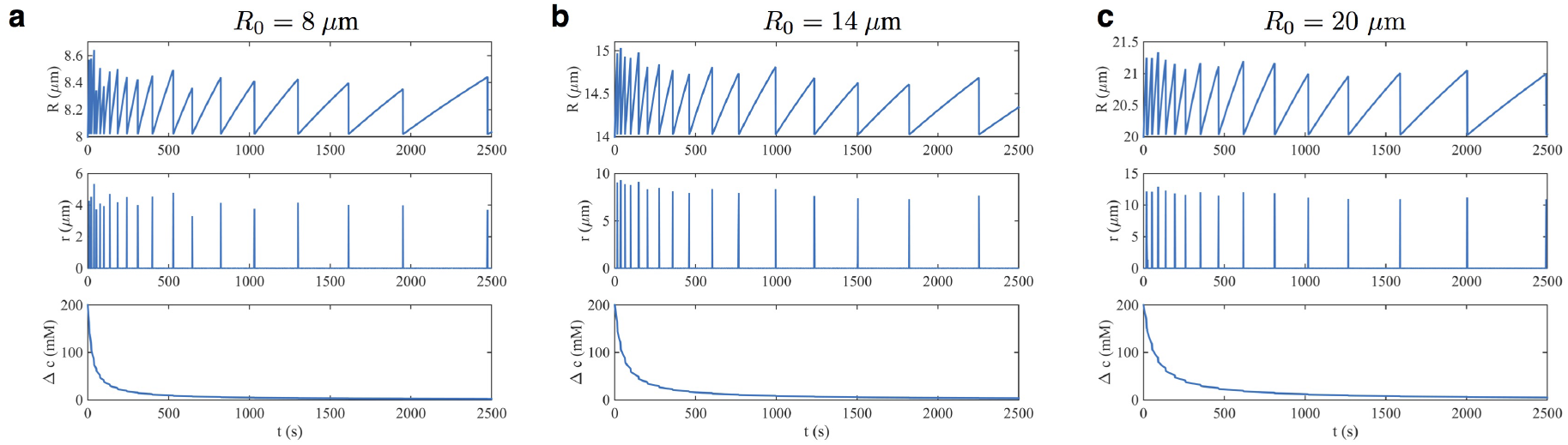
Influence of resting radius on the GUV swell-burst dynamics. All parameters are as shown in Table 1, except in panels a and c where *R*_0_ is set as indicated.

**Supplementary Figure 4:**
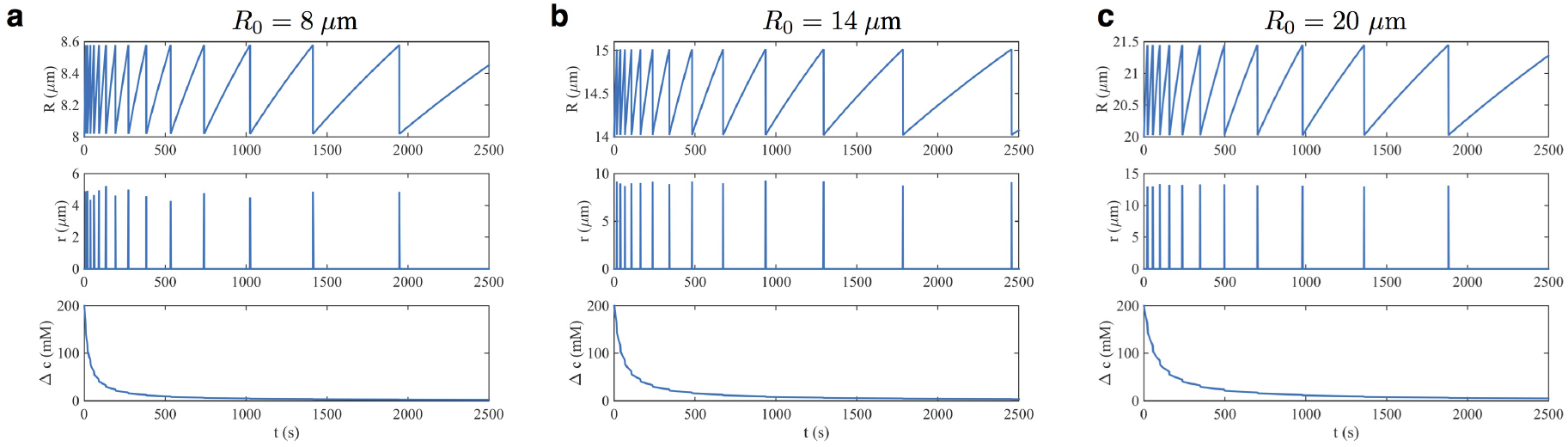
Solution to the deterministic model. All parameters are as shown in Table 1 except the pore opening strain set to 15%.

**Supplementary Figure 5:**
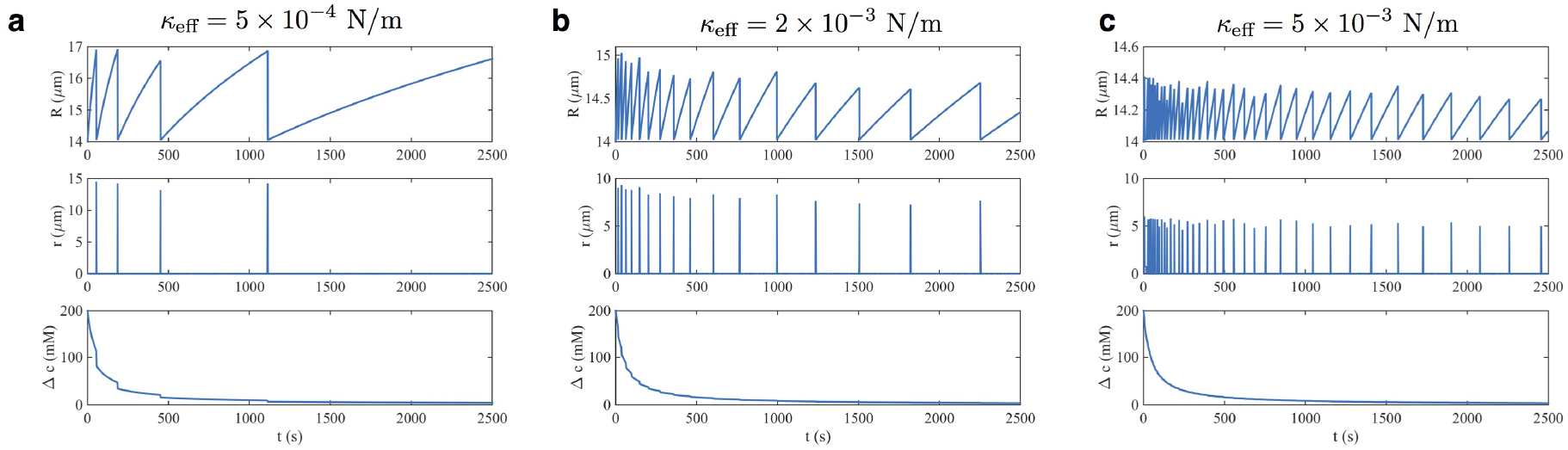
Influence of the effective stretching modulus on the GUV swell-burst dynamics. All parameters are as shown in Table 1, except in panels **a** and **c** where *κ*_eff_ is set as indicated, and *R*_0_ = 14 *μm*.

**Supplementary Figure 6:**
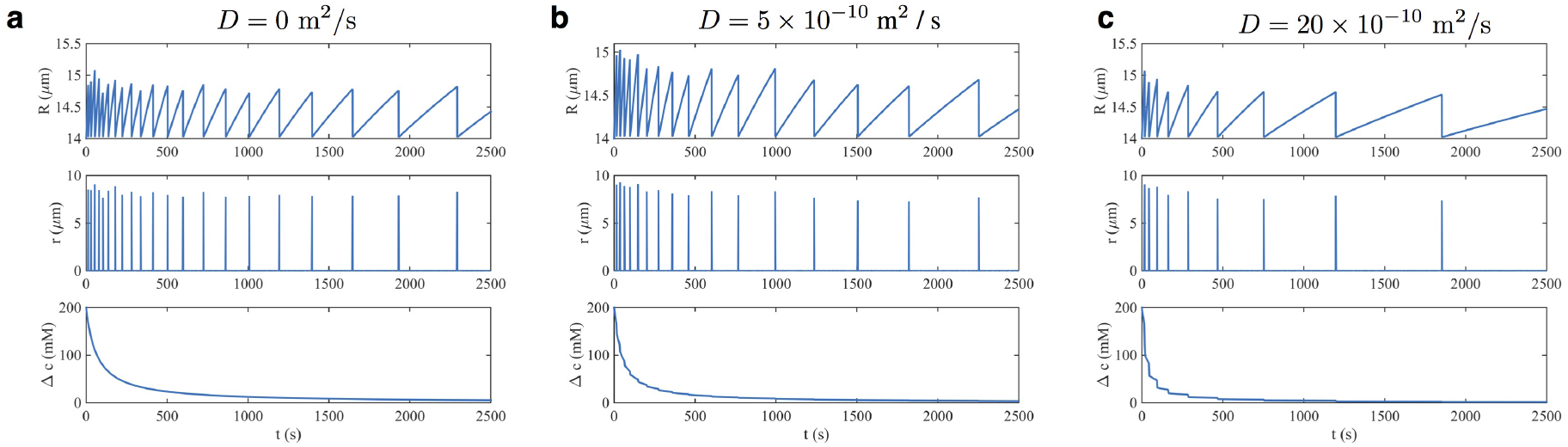
Influence of solute diffusion on the GUV swell-burst dynamics. All parameters are as shown in Table 1, except in panels **a** and **c** where *D* is set as indicated, and *R*_0_ = 14 *μm*.

**Supplementary Figure 7:**
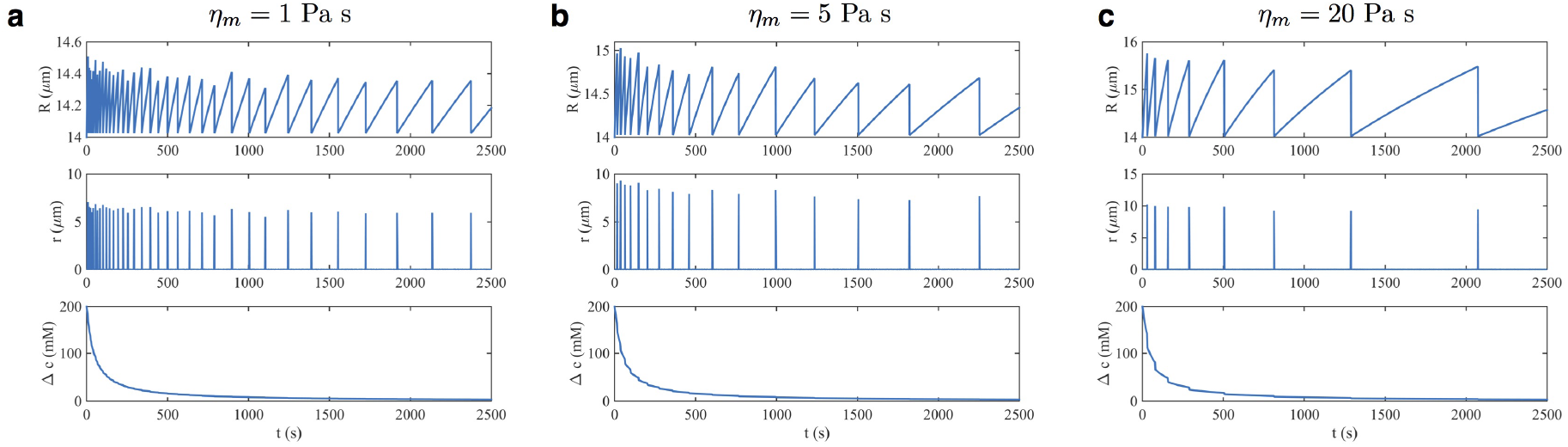
Influence of membrane viscosity on the GUV swell-burst dynamics. All parameters are as shown in Table 1, except in panels **a** and **c** where *η*_*m*_ is set as indicated, and *R*_0_ = 14 *μm*.

**Supplementary Figure 8:**
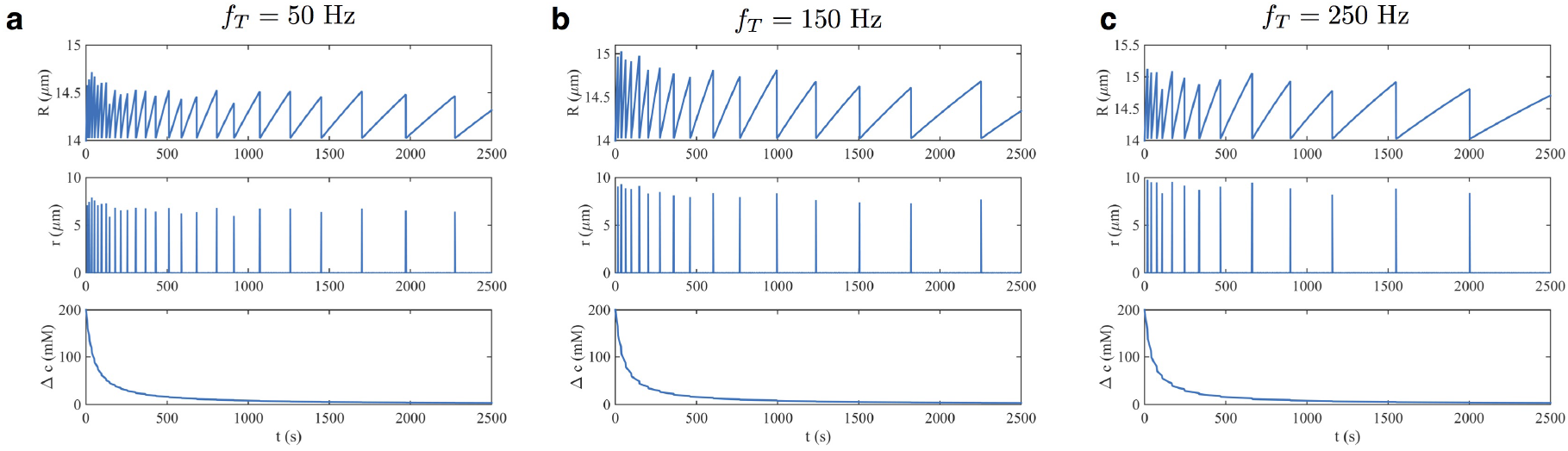
Influence of fluctuation frequency on the GUV swell-burst dynamics. All parameters are as shown in Table 1, except in panels **a** and **c** where *f*_*T*_ is set as indicated, and *R*_0_ = 14 *μm*.

**Supplementary Figure 9:**
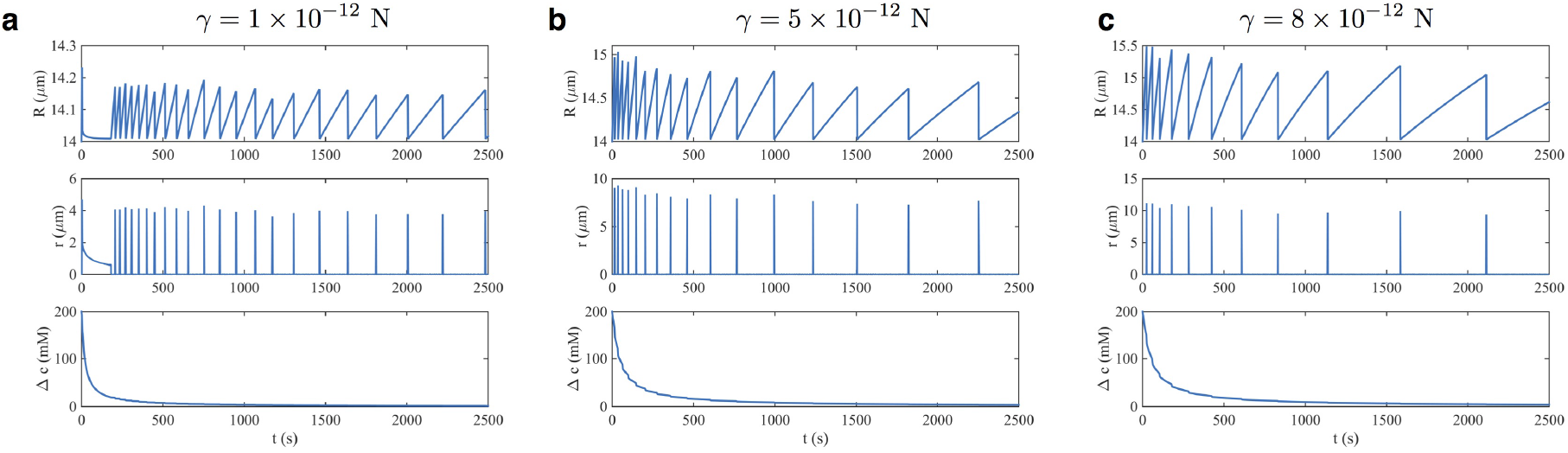
Influence of membrane line tension on the GUV swell-burst dynamics. All parameters are as shown in Table 1, except in panels **a** and **c** where *γ* is set as indicated, and *R*_0_ = 14 *μm*.

**Supplementary Figure 10:**
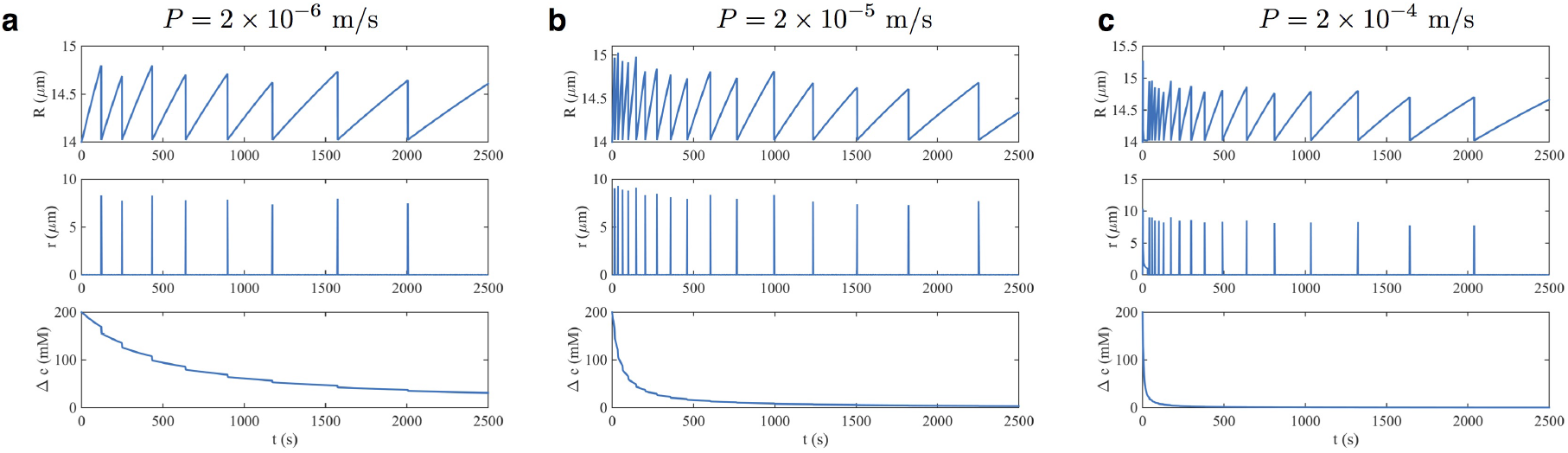
Influence of membrane solvent permeability on the GUV swell-burst dynamics. All parameters are as shown in Table 1, except in panels **a** and **c** where *P* is set as indicated, and *R*_0_ = 14 *μm*.

**Supplementary Figure 11:**
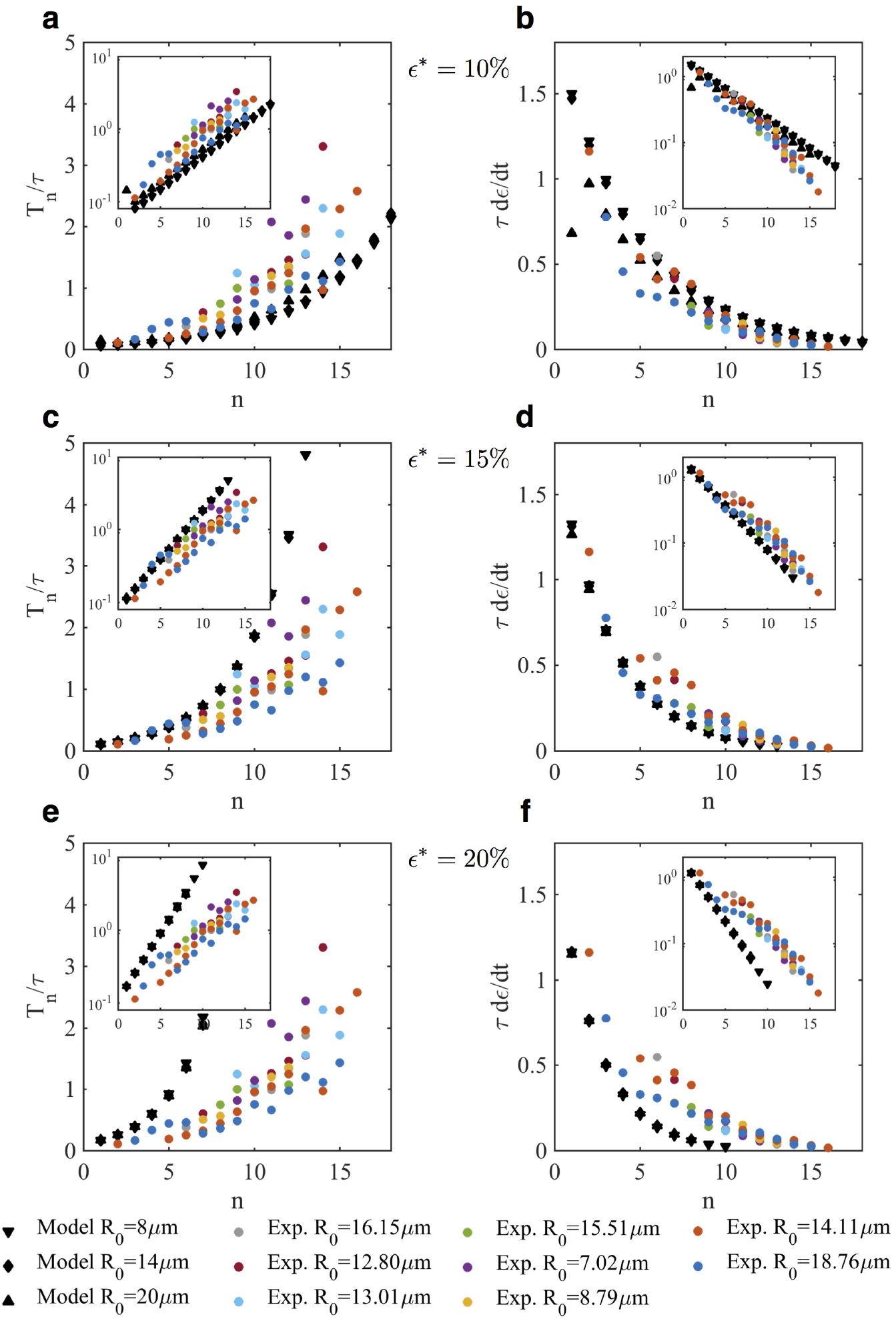
The deterministic model (constant strain to rupture *ε*\*) fails to match the experimental cycle period and strain rate. **a, b,** *ε*\* = 10%, **c, d,** *ε*\* = 15%, **e, f,** *ε*\* = 20%.

## Footnotes

1 http://rsb.info.nih.gov/ij/plugins/contour-plotter.html

## References

B. Alberts, A. Johnson, J. Lewis, D. Morgan, M. Raff, K. Roberts, and P. Walter. Molecular Biology of the Cell. Garland Science, 6 edition edition, Nov. 2014.

M. I. Angelova, S. Soléau, P. Méléard, F. Faucon, and P. Bothorel. Preparation of giant vesicles by external AC electric fields. Kinetics and applications. In C. Helm, M. Lösche, and H. Möhwald, editors, Trends in Colloid and Interface Science VI, number 89 in Progress in Colloid & Polymer Science, pages 127–131. Steinkopff, 1992. ISBN 978-3-7985-0913-9 978-3-7985-1680-9.

C. A. Aubin and R. J. Ryham. Stokes flow for a shrinking pore. Journal of Fluid Mechanics, 788:228–245, Feb. 2016. ISSN 1469-7645. doi: 10.1017/jfm.2015.699.

C. Berrier, M. Besnard, B. Ajouz, A. Coulombe, and A. Ghazi. Multiple mechanosensitive ion channels from escherichia coli, activated at different thresholds of applied pressure. The Journal of membrane biology, 151(2):175–187, 1996.

P. Blount, M. J. Schroeder, and C. Kung. Mutations in a bacterial mechanosensitive channel change the cellular response to osmotic stress. Journal of Biological Chemistry, 272(51):32150–32157, 1997.

F. Brochard, P. De Gennes, and P. Pfeuty. Surface tension and deformations of membrane structures: relation to two-dimensional phase transitions. Journal de Physique, 37(10):1099–1104, 1976.

F. Brochard-Wyart, P. G. de Gennes, and O. Sandre. Transient pores in stretched vesicles: role of leak-out. Physica A: Statistical Mechanics and its Applications, 278(1–2):32–51, Apr. 2000. ISSN 0378-4371. doi: 10.1016/S0378-4371(99)00559-2.

O. Christensen. Mediation of cell volume regulation by ca2+ influx through stretch-activated channels. Nature, 330:66–68, Nov. 1987.

E. R. Davies. Computer and Machine Vision: Theory, Algorithms, Practicalities. Academic Press, Apr. 2012. ISBN 978-0-12-386991-3.

D.W. Deamer and J. Bramhall. Special Issue: LiposomesPermeability of lipid bilayers to water and ionic solutes. Chemistry and Physics of Lipids, 40(2):167–188, June 1986. ISSN 0009-3084. doi: 10.1016/0009-3084(86)90069-1.

A. Diz-Muñoz, D. A. Fletcher, and O. D. Weiner. Use the force: membrane tension as an organizer of cell shape and motility. Trends in Cell Biology, 23(2):47–53, Feb. 2013. ISSN 0962-8924. doi: 10.1016/j.tcb.2012.09.006.

M. B. Elowitz, A. J. Levine, E. D. Siggia, and P. S. Swain. Stochastic gene expression in a single cell. Science, 297(5584):1183–1186, 2002.

A. Ertel, A. G. Marangoni, J. Marsh, F. R. Hallett, and J. M. Wood. Mechanical properties of vesicles. I. Coordinated analysis of osmotic swelling and lysis. Biophysical Journal, 64(2):426–434, Feb. 1993. ISSN 0006-3495. doi: 10.1016/S0006-3495(93)81383-3.

E. Evans, V. Heinrich, F. Ludwig, and W. Rawicz. Dynamic Tension Spectroscopy and Strength of Biomembranes. Biophysical Journal, 85(4):2342–2350, Oct. 2003. ISSN 0006-3495. doi: 10.1016/S0006-3495(03)74658-X.

R. Fettiplace and D. A. Haydon. Water permeability of lipid membranes. Physiological Reviews, 60(2):510–550, Apr. 1980.

F. R. Hallett, J. Marsh, B. G. Nickel, and J. M. Wood. Mechanical properties of vesicles. II. A model for osmotic swelling and lysis. Biophysical Journal, 64(2):435–442, Feb. 1993. ISSN 0006-3495. doi: 10.1016/S0006-3495(93)81384-5.

J. Happel and H. Brenner. Low Reynolds Number Hydrodynamics: With Special Applications to Particulate Media. Springer Science & Business Media, Sept. 1983. ISBN 978-90-247-2877-0.

W. Helfrich and R.-M. Servuss. Undulations, steric interaction and cohesion of fluid membranes. Il Nuovo Cimento D, 3(1):137–151, Jan. 1984. ISSN 0392-6737. doi: 10.1007/BF02452208.

J. C. S. Ho, P. Rangamani, B. Liedberg, and A. N. Parikh. Mixing Water, Transducing Energy, and Shaping Membranes: Autonomously Self-Regulating Giant Vesicles. Langmuir, Feb. 2016. ISSN 0743-7463. doi: 10.1021/acs.langmuir.5b04470.

E. K. Hoffmann, I. H. Lambert, and S. F. Pedersen. Physiology of Cell Volume Regulation in Vertebrates. Physiological Reviews, 89(1):193–277, Jan. 2009. ISSN 0031-9333, 1522-1210. doi: 10.1152/physrev. 00037.2007.

T. T. Hormel, S. Q. Kurihara, M. K. Brennan, M. C. Wozniak, and R. Parthasarathy. Measuring Lipid Membrane Viscosity Using Rotational and Translational Probe Diffusion. Physical Review Letters, 112(18):188101, May 2014. doi: 10.1103/PhysRevLett.112.188101.

M. A. Idiart and Y. Levin. Rupture of a liposomal vesicle. Physical Review E, 69(6):06 1922., June 2004. doi: 10.1103/PhysRevE.69.061922.

F. Jülicher, A. Ajdari, and J. Prost. Modeling molecular motors. Reviews of Modern Physics, 69(4):1269, 1997.

E. Karatekin, O. Sandre, H. Guitouni, N. Borghi, P.-H. Puech, and F. Brochard-Wyart. Cascades of Transient Pores in Giant Vesicles: Line Tension and Transport. Biophysical Journal, 84(3):1734–1749, Mar. 2003. ISSN 0006-3495. doi: 10.1016/S0006-3495(03)74981-9.

M. M. Koslov and V. S. Markin. A theory of osmotic lysis of lipid vesicles. Journal of Theoretical Biology, 109(1):17–39, July 1984. ISSN 0022-5193. doi: 10.1016/S0022-5193(84)80108-3.

R. Kubo. The fluctuation-dissipation theorem. Reports on progress in physics, 29(1):255, 1966.

N. Levina, S. Tötemeyer, N. R. Stokes, P. Louis, M. A. Jones, and I. R. Booth. Protection of escherichia coli cells against extreme turgor by activation of mscs and mscl mechanosensitive channels: identification of genes required for mscs activity. The EMBO journal, 18(7):1730–1737, 1999.

P. W. Linder, L. R. Nassimbeni, A. Polson, and A. L. Rodgers. The diffusion coefficient of sucrose in water. A physical chemistry experiment. Journal of Chemical Education, 53(5):330, May 1976. ISSN 0021-9584. doi: 10.1021/ed053p330.

J. D. Litster. Stability of lipid bilayers and red blood cell membranes. Physics Letters A, 53(3):193–194, June 1975. ISSN 0375-9601. doi: 10.1016/0375-9601(75)90402-8.

K. Oglecka, P. Rangamani, B. Liedberg, R. S. Kraut, and A. N. Parikh. Oscillatory phase separation in giant lipid vesicles induced by transmembrane osmotic differentials. eLife, 3:e03695, Oct. 2014. ISSN 2050-084X. doi: 10.7554/eLife.03695.

K. Olbrich, W. Rawicz, D. Needham, and E. Evans. Water permeability and mechanical strength of polyunsaturated lipid bilayers. Biophysical Journal, 79(1):321–327, July 2000. ISSN 0006-3495.

G. Oster. Brownian ratchets: Darwin’s motors. Nature, 417(6884):25–25, May 2002. ISSN 0028-0836. doi: 10.1038/417025a.

P. Peterlin and V. Arrigler. Electroformation in a flow chamber with solution exchange as a means of preparation of flaccid giant vesicles. Colloids and Surfaces B: Biointerfaces, 64(1):77–87, June 2008. ISSN 0927-7765. doi: 10.1016/j.colsurfb.2008.01.004.

P. Peterlin, V. Arrigler, H. Diamant, and E. Haleva. Permeability of Phospholipid Membrane for Small Polar Molecules Determined from Osmotic Swelling of Giant Phospholipid Vesicles. In A. Iglič, editor, Advances in Planar Lipid Bilayers and Liposomes, volume 16, pages 301–335. Academic Press, 2012a.

P. Peterlin, V. Arrigler, E. Haleva, and H. Diamant. Law of corresponding states for osmotic swelling of vesicles. Soft Matter, 8(7):2185, 2012b. ISSN 1744-683X, 1744-6848. doi: 10.1039/c1sm06670f.

D. Popescu and A. G. Popescu. The working of a pulsatory liposome. Journal of Theoretical Biology, 254 (3):515–519, Oct. 2008. ISSN 0022-5193. doi: 10.1016/j.jtbi.2008.07.009.

C. A. M. L. Porta, A. Ghilardi, M. Pasini, L. Laurson, M. J. Alava, S. Zapperi, and M. B. Amar. Osmotic stress affects functional properties of human melanoma cell lines. The European Physical Journal Plus, 130(4):1–15, Apr. 2015. ISSN 2190-5444. doi: 10.1140/epjp/i2015-15064-x.

T. Portet and R. Dimova. A New Method for Measuring Edge Tensions and Stability of Lipid Bilayers: Effect of Membrane Composition. Biophysical Journal, 99(10):3264–3273, Nov. 2010. ISSN 0006-3495. doi: 10.1016/j.bpj.2010.09.032.

R. P. Rand. Probing the role of water in protein conformation and function. Philosophical Transactions of the Royal Society of London B: Biological Sciences, 359(1448):1277–1285, Aug. 2004. ISSN 0962-8436, 1471-2970. doi: 10.1098/rstb.2004.1504.

P. Rangamani, K. K. Mandadap, and G. Oster. Protein-induced membrane curvature alters local membrane tension. Biophysical journal, 107(3):751–762, 2014.

R. Ryham, I. Berezovik, and F. S. Cohen. Aqueous Viscosity Is the Primary Source of Friction in Li-pidic Pore Dynamics. Biophysical Journal, 101(12):2929–2938, Dec. 2011. ISSN 0006-3495. doi: 10.1016/j.bpj.2011.11.009.

C. A. Schneider, W. S. Rasband, K. W. Eliceiri, et al. Nih image to imagej: 25 years of image analysis. Nat methods, 9(7):671–675, 2012.

U. Seifert. Stochastic thermodynamics: principles and perspectives. The European Physical Journal B, 64(3-4):423–431, 2008.

K. M. Stroka, H. Jiang, S.-H. Chen, Z. Tong, D.Wirtz, S. X. Sun, and K. Konstantopoulos. Water Permeation Drives Tumor Cell Migration in Confined Microenvironments. Cell, 157(3):611–623, Apr. 2014. ISSN 0092-8674. doi: 10.1016/j.cell.2014.02.052.

H. Turlier, D. Fedosov, B. Audoly, T. Auth, N. Gov, C. Sykes, J.-F. Joanny, G. Gompper, and T. Betz. Equilibrium physics breakdown reveals the active nature of red blood cell flickering. Nature Physics, 2016.

J.M. Wood. Osmosensing by Bacteria: Signals and Membrane-Based Sensors. Microbiology and Molecular Biology Reviews, 63(1):230–262, Jan. 1999. ISSN 1092-2172, 1098-5557.

